# A combinatorial anticancer drug screen identifies off-target effects of epigenetic chemical probes

**DOI:** 10.1101/2022.04.14.488411

**Authors:** Samir H. Barghout, Mandeep K. Mann, Yifan Yu, Aaron D. Schimmer, Matthieu Schapira, Cheryl H. Arrowsmith, Dalia Barsyte-Lovejoy

**Affiliations:** Structural Genomics Consortium, University of Toronto, Toronto, ON, Canada; Department of Pharmacology & Toxicology, Faculty of Medicine, University of Toronto, Toronto, ON, Canada; Department of Pharmacology & Toxicology, Faculty of Pharmacy, Tanta University, Tanta, Egypt; Department of Medical Biophysics, Faculty of Medicine, University of Toronto, Toronto, ON, Canada; Princess Margaret Cancer Centre, University Health Network, Toronto, ON, Canada

## Abstract

Anticancer drug response is determined by genetic and epigenetic mechanisms. To identify the epigenetic regulators of anticancer drug response, we conducted a chemical epigenetics screen using chemical probes that target different epigenetic modulators. In this screen, we tested 31 epigenetic probes in combination with 14 mechanistically diverse anticancer agents and identified 8 epigenetic probes that significantly potentiate the cytotoxicity of TAK-243, a first-in-class ubiquitin-activating enzyme (UBA1) inhibitor evaluated in several solid and hematologic malignancies. These probes are TP-472, GSK-864, A-196, UNC1999, SGC-CBP30 and PFI-4 (and its related analogs GSK6853 and GSK5959), and they target BRD9/7, mutant IDH1, SUV420H1/2, EZH2/1, p300/CBP and BRPF1B, respectively. In contrast to epigenetic probes, negative control compounds did not have a significant impact on TAK-243 cytotoxicity. Potentiation of TAK-243 cytotoxicity was associated with reduced ubiquitylation and induction of apoptosis. Mechanistically, epigenetic probes exerted their potentiation by inhibiting the efflux transporter ABCG2 without inducing significant changes in the ubiquitylation pathways or ABCG2 expression levels. The identified probes shared chemical scaffold similarities with TAK-243 and could potentially interact with ABCG2 as assessed by docking analysis. Based on these data, we have developed a cell-based assay that can quantitatively evaluate ABCG2 inhibition by drug candidates. In conclusion, our study identifies epigenetic chemical probes that profoundly potentiate TAK-243 cytotoxicity through off-target ABCG2 inhibition. We also provide experimental evidence that several negative control compounds cannot exclude a subset of off-target effects of chemical probes. Finally, potentiation of TAK-243 cytotoxicity can serve as a quantitative measure of ABCG2-inhibitory activity.

## Introduction

Anticancer drug response is determined by numerous factors including the expression levels of the drug targets and/or transporters, metabolic alterations, activity of pro-survival and death signaling pathways, and cell-extrinsic factors related to the tumor microenvironment^1, 2^. These factors are regulated by genetic and/or epigenetic mechanisms^3, 4^. While genetic regulators have been extensively studied in numerous contexts, epigenetic determinants of anticancer drug response are less understood^5^.

In this respect, several epigenetic mechanisms have been implicated in regulating the response to cytotoxic and targeted therapies in numerous malignancies^4, 6–11^. Accordingly, a number of combination therapies incorporating epigenetic drugs have been proposed to improve the anticancer drug response^5, 12^. Nonetheless, a systematic understanding of the role of different epigenetic regulators in the response to anticancer drug therapy is still lacking^5^.

A number of genetic screens focused on epigenetic targets have been conducted to identify the epigenetic regulators of tumor progression^13–16^, with few screens performed in the context of anticancer drug therapy. The use of these genetic approaches to screen for synthetic lethal interactions with multiple anticancer drugs may be technically challenging. Thus, the use of a chemical epigenetics approach may be a more viable strategy to allow rapid identification of drug combinations that can be fast-tracked or optimized for use as anticancer therapeutic modalities^17^. Endeavors of the Structural Genomics Consortium (SGC) and other members of the scientific community have culminated in the development of a chemical biology toolbox comprising collections of chemical probes that target epigenetic modulators from diverse families in a potent and selective manner^18–20^.

Here, we leveraged the SGC toolbox of epigenetic probes and performed a combinatorial screen with a panel of anticancer agents to gain novel insights into the chemical biology of these probes and help identify more effective drug combinations that may achieve better therapeutic outcomes. We identified 8 epigenetic chemical probes targeting different epigenetic modulators that profoundly potentiate the cytotoxicity of the ubiquitin-activating enzyme (UBA1) inhibitor TAK-243 in cell lines of different origin. Compared to other anticancer agents, the potentiation was relatively selective for TAK-243. Through functional and mechanistic experiments, we demonstrated the potentiation was mediated by an off-target inhibition of the efflux ABC transporter ABCG2 which, unexpectedly, was not observed with negative epigenetic control compounds. Additionally, we exploited this profound potentiation to develop a cellular assay for the quantitative assessment of ABCG2 inhibition by drug candidates.

## Materials and methods

### Chemicals and reagents

Epigenetic probes described in **Table S1** were obtained from the Structural Genomics Consortium (Toronto, Canada). TAK-243 was purchased from Active Biochem (catalog# A-1384), pevonedistat (MLN4924/TAK-924) (catalog #15217), TAK-981 (catalog# 32741), bortezomib (catalog# 10008822), mitoxantrone (catalog# 14842), panobinostat (catalog# 13280), lenalidomide (catalog# 14643), imatinib (catalog# 13139), vincristine (catalog# 11764), cyclophosphamide (catalog# 13849), and cisplatin (catalog# 13119) from Cayman chemicals, zosuquidar (catalog# 5456) and Ko143 (catalog# 3241) from Tocris, GSK-5959 (catalog# HY-18665), inobrodib (catalog# HY-111784), and ivosidenib (catalog# HY-18767) from MedChemExpress. Cisplatin was dissolved in molecular-grade water and all other drugs were dissolved in DMSO.

### Cell culture

OCI-AML2, K562, and MV4-11 were cultured in Iscove’s modified Dulbecco’s medium (IMDM), NB4, U937, MDAY-D2, KMS-11, RPMI 8226 and Jurkat cells in Roswell Park Memorial Institute (RPMI) medium, and HEK293, A549, Hep G2 and MCF7 cells in Dulbecco’s Modified Eagle Medium (DMEM). All culture media were supplemented with 10% fetal bovine serum (FBS) and 1% penicillin/streptomycin. Cell lines were incubated at 37^°^C, 5% CO_2_ and 95% humidity. OCI-AML2, OCI-M2 and MDAY-D2 cells were obtained as a gift from Dr. Mark Minden (University Health Network, Toronto, Canada). Other cell lines were obtained from the American Type Culture Collection (ATCC) organization.

### Combinatorial chemical screen

A549 cells were seeded in 96-well plates (5,000 cells/well) and left to adhere overnight. Cells were subsequently treated with epigenetic probes alone and in combination with anticancer agents at single concentrations as described in **Tables S1 and S2** for 72h. Concentrations that did not significantly affect cell viability were selected to allow for capturing potentiation/synergism. Each epigenetic probe was represented by two technical replicates and data were obtained from 2-3 independent experiments. DMSO and anticancer drug controls were randomly distributed in different locations in the plates to account for any location-dependent variabilities in cell growth. Viability was measured after 72h using the Resazurin assay. Percent viability was obtained by normalization to DMSO-treated cells in plates treated with epigenetic probes alone, and to anticancer drug-treated cells in plates treated with combinations to account for reductions in viability elicited by anticancer agents. The proliferation of cells over 72h was also monitored using phase contrast imaging and subsequent analysis by Incucyte Software.

### Cytotoxicity assays

CellTiter 96^®^ AQueous MTS Reagent Powder was purchased from Promega (catalog# G1111) and the MTS reagent was prepared as per the manufacturer’s instructions. Resazurin sodium salt (Alamar blue) was purchased from Sigma-Aldrich (catalog# R7017) and dissolved in water at a concentration of 0.04 mg/mL. Cells were counted and plated in 96-well plates at appropriate densities: OCI-AML2 (25,000/well), OCI-M2 (25,000/well), K562 (10,000/well), MV4-11 (25,000/well), RPMI 8226 (25,000/well), KMS-11 (25,000/well), NB4 (25,000/well), U937 (10,000/well), HepG2 (5,000/well), MDAY-D2 (10,000/well), HEK293 (5,000/well) and Jurkat (10,000/well). Adherent cells were allowed to adhere overnight. Cells were subsequently treated with increasing concentrations of the drug(s) under investigation. After 72h of incubation, the MTS or resazurin solutions were directly added to the media at a ratio of 1:5. For the MTS assay, the absorbance was measured at 490 nm using SpectraMax microplate reader (Molecular Devices, USA). For the resazurin assay, the fluorescence was measured at 560_Ex_/590_Em_ using CLARIOstar microplate reader (BMG LABTECH, Mandel Scientific, Canada). Viability was then calculated as a percentage of control-treated cells and concentration-response curves were constructed. The IC_50_ values were calculated using the non-linear regression function in GraphPad Prism (Version 9.0.0, GraphPad Software Inc, LLC).

### Cellular thermal shift assay (CETSA)

The CETSA assay was performed as previously described^21^. In brief, cells were pre-treated with epigenetic probes for 4h and 48h. Untreated and probe-treated cells were then treated with TAK-243 for 1h. Cells were subsequently washed with phosphate-buffered saline (PBS) and re-suspended in PBS containing protease inhibitors (Roche, Canada). Cells were heated at 54^°^C for 3 min in a thermal cycler (VeritiPro, Applied Biosystems). This temperature was previously selected as the temperature corresponding to the maximal thermal shift of UBA1^22^. Cells were lysed by 3 freeze-thaw-vortex cycles using liquid nitrogen and a thermal cycler set at RT. Lysates were subsequently centrifuged at 20,000 g for 20 min and supernatants were collected and stored at −20^°^C until immunoblotting.

### Immunoblotting

Whole cell lysates were prepared by washing cells with PBS (pH=7.4) followed by lysis in a buffer composed of 20 mM Tris-HCl pH 8, 150 mM NaCl, 1 mM EDTA, 10 mM MgCl_2_, 0.5% Triton X-100, 12.5 U/mL Benzonase^®^ nuclease (catalog#E8263; Sigma-Aldrich, Canada), and complete EDTA-free protease inhibitor cocktail (Roche, Canada). Total protein was quantified using Pierce™ BCA protein assay kit (Thermo Fisher Scientific, Canada). Samples were subsequently denatured by boiling at 95°C for 5 min, loaded in equal amounts, and fractionated by NuPAGE™ 4-12% gradient gels (Thermo Fisher Scientific, Canada) using sodium dodecyl sulphate-polyacrylamide gel electrophoresis (SDS-PAGE). Fractionated proteins were subsequently transferred to polyvinylidene difluoride (PVDF) membranes and probed using appropriate primary antibodies (**Table S3**). Secondary goat-anti rabbit-IR800 (catalog# 926-32211) and donkey anti-mouse-IR680 (catalog#926-68072) were purchased from LI-COR Biosciences.

### shRNA-mediated knockdown experiments

shRNAs targeting *ABCG2* were obtained from Sigma-Aldrich (product# SHCLNG-NM_004827). Viral supernatants were prepared by transfecting 293T cells with *GFP*- or *ABCG-*targeting shRNAs encoded in pLKO.1 vectors using X-tremeGENE 360 transfection reagent (Roche, Germany) as per the manufacturer’s instructions. To transduce A549 and RPMI 8226 cells, viral supernatants (0.5 mL) of *GFP*- and *ABCG2*-targeting shRNAs and polybrene (8 µg/mL) were added to cells (200K and 1000K cells, respectively) for 24h. Media were subsequently replaced, and cells selected with puromycin (1 µg /mL) for 72h. Cell lysates were prepared, and knockdown confirmed by immunoblotting. Sequences of *ABCG2*-targeting shRNAs are: sh*ABCG2*#1: CCT GCC AAT TTC AAA TGT AAT; sh*ABCG2*#2: TAA CAT CTG CTA TCG AGT AAA.

### Docking analysis

Ligprep (Schrodinger) was used to prepare the ligands using default settings. The missing side chain atoms of the cryo-EM structures of ABCG2 (PDB: 6ETI^23^, 7NEQ^24^, 70J8^25^) were added with ICM-Pro^26^ (Molsoft) followed by preparation with PrepWizard^27^ (Schrodinger) for the assignment of the bond order, assessment of correct protonation states, and optimization of the hydrogen bond assignment at pH 7.3, and restrained minimization using the OPLS4 force field. Receptor grids were calculated at the centroid of the crystallized ligand for each respective complex. The library was docked to the structures using Glide SP^28^ (Schrodinger, New York). The top-scoring pose of each ligand was chosen for final analysis and visualization.

### Data analysis

Median inhibitory concentration (IC_50_) values were calculated using the nonlinear regression function in GraphPad Prism software version 9.0.0 (GraphPad Software Inc.). Chemical structures were drawn using ChemDraw version 20.1.1 (PerkinElmer Inc., USA).

## Results

### A combinatorial screen identifies hits that profoundly potentiate TAK-243 cytotoxicity

The SGC has developed a chemical biology toolbox comprising chemical probes that target epigenetic modulators in a potent and selective manner^19, 20^. Leveraging these compounds, we performed a combinatorial screen with a panel of anticancer agents to identify probes that potentiate the cytotoxicity of these drugs. Specifically, we treated A549 lung adenocarcinoma cells with 31 epigenetic probes alone and in combination with 14 structurally and mechanistically distinct anticancer agents. To facilitate capturing potentiation/synergism, we used epigenetic probes and anticancer agents at concentrations that do not cause significant cytotoxicity (**Table S1** and **S2**). We subsequently assessed the impact of these compounds on short-term proliferation and viability at 72h.

As measured by the Resazurin assay, most epigenetic probes and anticancer agents did not induce significant cytotoxicity alone nor in combination (**Fig. 1A and S1)**. Notably, however, 6 epigenetic probes (TP-472, GSK-864, A-196, UNC1999, SGC-CBP30, and PFI-4) significantly potentiated TAK-243 cytotoxicity (**Fig. 1A and S1A)**. TAK-243 is a first-in-class inhibitor of the ubiquitin-activating enzyme UBA1 that initiates the ubiquitylation cascade^22, 29, 30^. As assessed by phase contrast imaging, combination of these epigenetic probes and TAK-243 abrogated cellular proliferation and viability as early as 24h after treatment, corroborating the Resazurin assay data (**Fig. 1B** and **S2)**. Of note, the identified probes target functionally distinct epigenetic modulators from different families (**Table S1**). Taken together, our screen identified mechanistically distinct epigenetic probes that elicit a highly significant potentiation of TAK-243.

**Figure 1.**
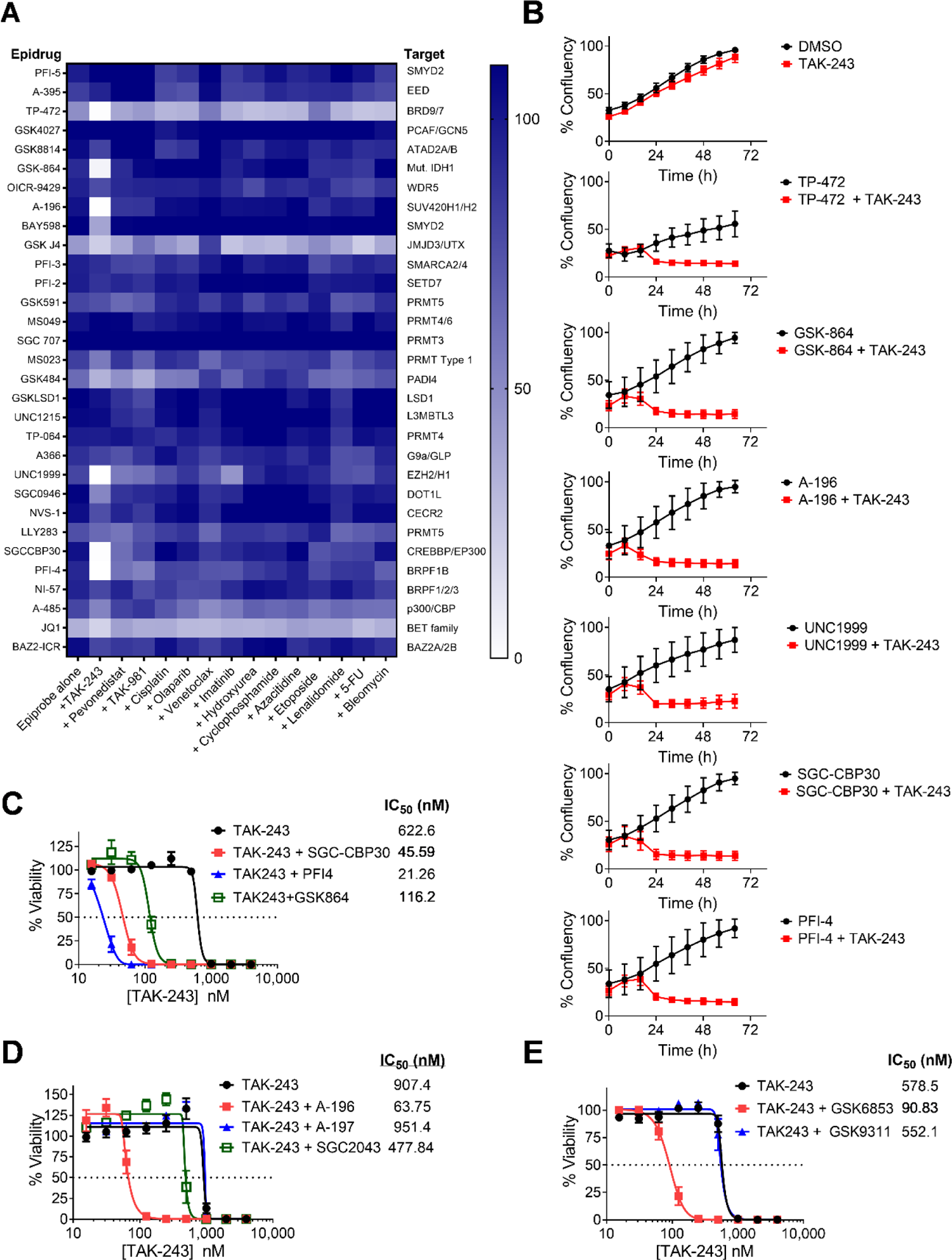
A combinatorial screen identifies epigenetic probes that profoundly potentiate TAK-243 cytotoxicity. A) A heat map showing 31 epigenetic probes used alone (at concentrations from 1-5 µM) or in combination with 14 anticancer agents in A549 lung adenocarcinoma cells. Targets of the epigenetic probes are shown B) Proliferation curves of A549 lung adenocarcinoma cells after treatment with DMSO, TAK-243 (100 nM), and epigenetic probes (5 µM) alone or in combination with TAK-243. Proliferation was represented as % confluency as measured by the Incucyte imaging software over 72h. Error bars represent SD of % confluency calculated from 8-10 images. C) Concentration-response curve of TAK-243 alone or in combination with SGC-CBP30, PFI-4, GSK-864 in RPMI-8226 cells. Insert: median inhibitory concentration (IC_50_) values D) Concentration-response curve of TAK-243 alone or in combination with A-196 (5 µM) or its negative controls A-197 and SGC-2043 (5 µM) in RPMI-8226 myeloma cells E) Concentration-response curve of TAK-243 alone or in combination with GSK6853 and its negative control GSK9311 in RPMI-8226 cells. In the screen and all validation experiments, viability was assessed using the Resazurin assay after treatment for 72h. In concentration-response curves, data points represent the mean ± SEM of at least 3 independent experiments.

### Screen hits potentiate TAK-243 cytotoxicity in cell lines of different origin

To validate the screen results, we tested the potentiation in different cell lines, using an additional viability assay, and over a wide range of TAK-243 concentrations. Specifically, we treated RPMI 8226 and KMS11 myeloma cells, known to be relatively insensitive to TAK-243^29, 31^, with increasing concentrations of TAK-243 alone and in combination with the hits identified from the screen and assessed viability by the Resazurin assay after 72h. In line with the screen results, combination with these epigenetic probes profoundly potentiated TAK-243 cytotoxicity with a 4- to 30-fold reduction in the IC_50_ of TAK-243 (**Fig. 1C-E and S3A)**.

Together with chemical probes, the SGC has developed structurally related, functionally inactive compounds to serve as negative controls that can be used to exclude off-target effects^32^. Thus, in parallel, we tested the effect of negative controls of A-196 (A-197 and SGC-2043), UNC1999 (UNC2400), and TP-472 (TP-472N) on TAK-243 cytotoxicity. We also tested the PFI-4 analog GSK6853 for which a negative control known as GSK9311 is available. With the exception of TP-472N, all other negative controls did not phenocopy their active probes with no significant changes in the IC_50_ of TAK-243 (**Fig. 1D-E and S3)**.

To expand the cell lines tested, we treated a panel of 14 cell lines of different origin with increasing concentrations of TAK-243 alone and in combination with A-196 and assessed viability by a distinct viability assay, the MTS assay, after treatment for 72h. Of the 14 cell lines tested, A-196 potentiated TAK-243 cytotoxicity in 10 cell lines with a 2- to 40-fold reduction in the IC_50_ of TAK-243 (**Fig. 2**). Taken together, these data confirm the screen results as the identified hits, but not their negative controls, potentiated TAK-243 cytotoxicity in cell lines of different origin using different viability assays.

**Figure 2.**
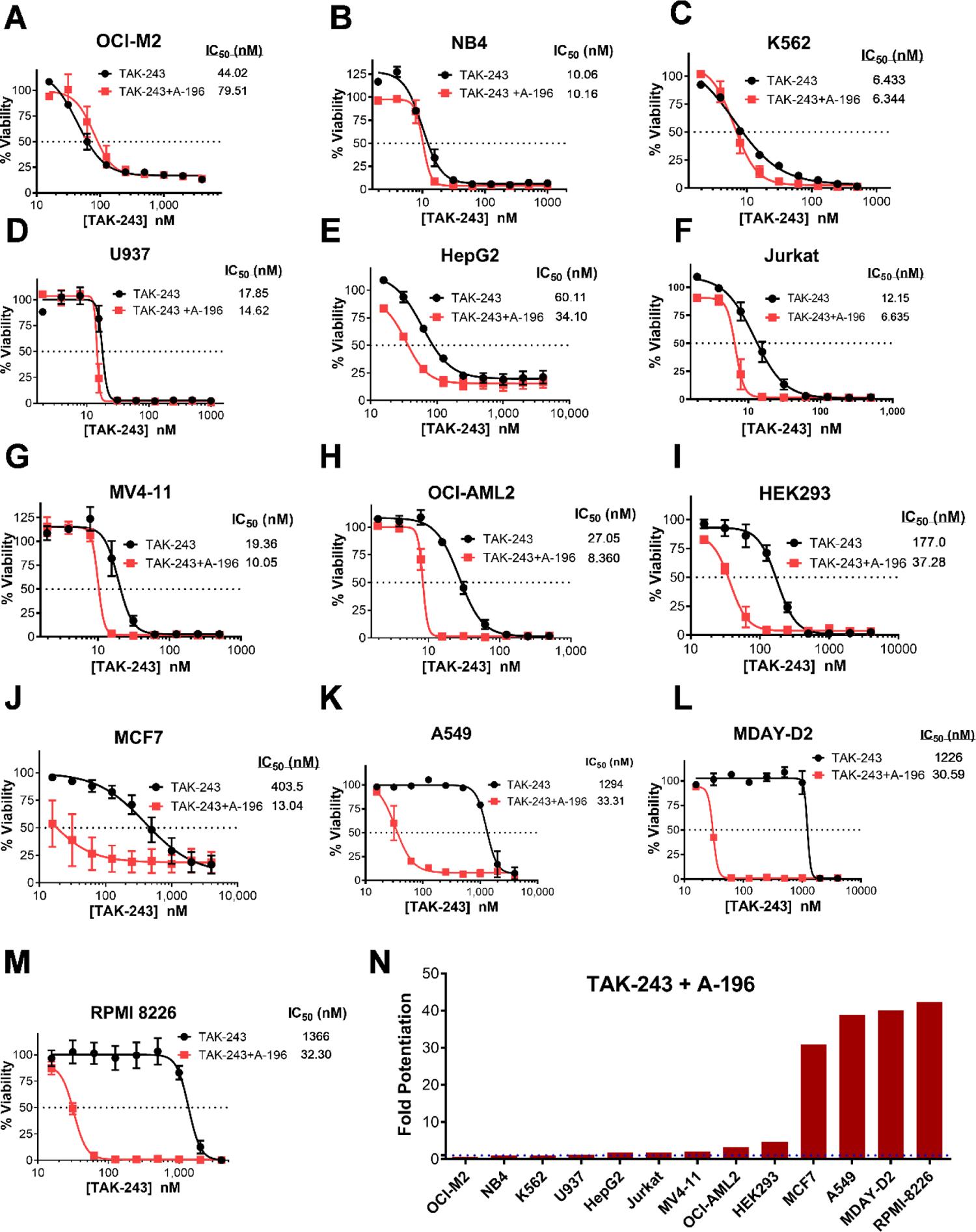
A-196 potentiates TAK-243 cytotoxicity in cell lines of different origin. A-M) Concentration-response curves of TAK-243 alone or in combination with A-196 (5 µM) in 13 cancer cell lines of different origin. Insert: median inhibitory concentration (IC_50_) values. All data points represent the mean ± SEM of 2-3 independent experiments N) Bar chart showing fold-potentiation of TAK-243 when combined with A-196 (5 µM) in different cell lines. Viability was assessed using the MTS assay after treatment for 72h.

### Potentiation of TAK-243 cytotoxicity is associated with reduced ubiquitylation and induction of apoptosis

TAK-243 selectively inhibits UBA1, the major cellular enzyme responsible for activating ubiquitylation^30^. Thus, TAK-243 treatment is anticipated to reduce the abundance of ubiquitylated proteins, leading to the induction of cell death^22, 29^. To assess the molecular changes associated with the potentiation of TAK-243 cytotoxicity, we pre-treated RPMI 8226 cells with the screen hits or their negative controls for 24h followed by TAK-243 treatment for 8h, and measured changes in the abundance of ubiquitylated histone H2A (Ub-H2A), Ub-H2B, and cleaved poly (ADP-ribose) polymerase (PARP) that serves as a marker of apoptosis^22, 33^. Treatment with vehicle, epigenetic probes, or TAK-243 alone did not induce significant changes in Ub-H2A, Ub-H2B or cleaved PARP (**Fig. 3**). In contrast, combinatorial treatment with the screen hits and TAK-243 resulted in profound reduction in the ubiquitylation of H2A and H2B, and a significant induction of PARP cleavage. Additionally, combinatorial treatment with the negative controls A-197 and GSK9311 did not cause significant changes in Ub-H2A, Ub-H2B, or PARP cleavage (**Fig. 3A**). Taken together, the cytotoxicity observed after combinatorial treatment with the screen hits and TAK-243 is associated with molecular changes suggestive of potentiation of TAK-243 action on ubiquitylation and downstream events leading to apoptosis.

**Figure 3.**
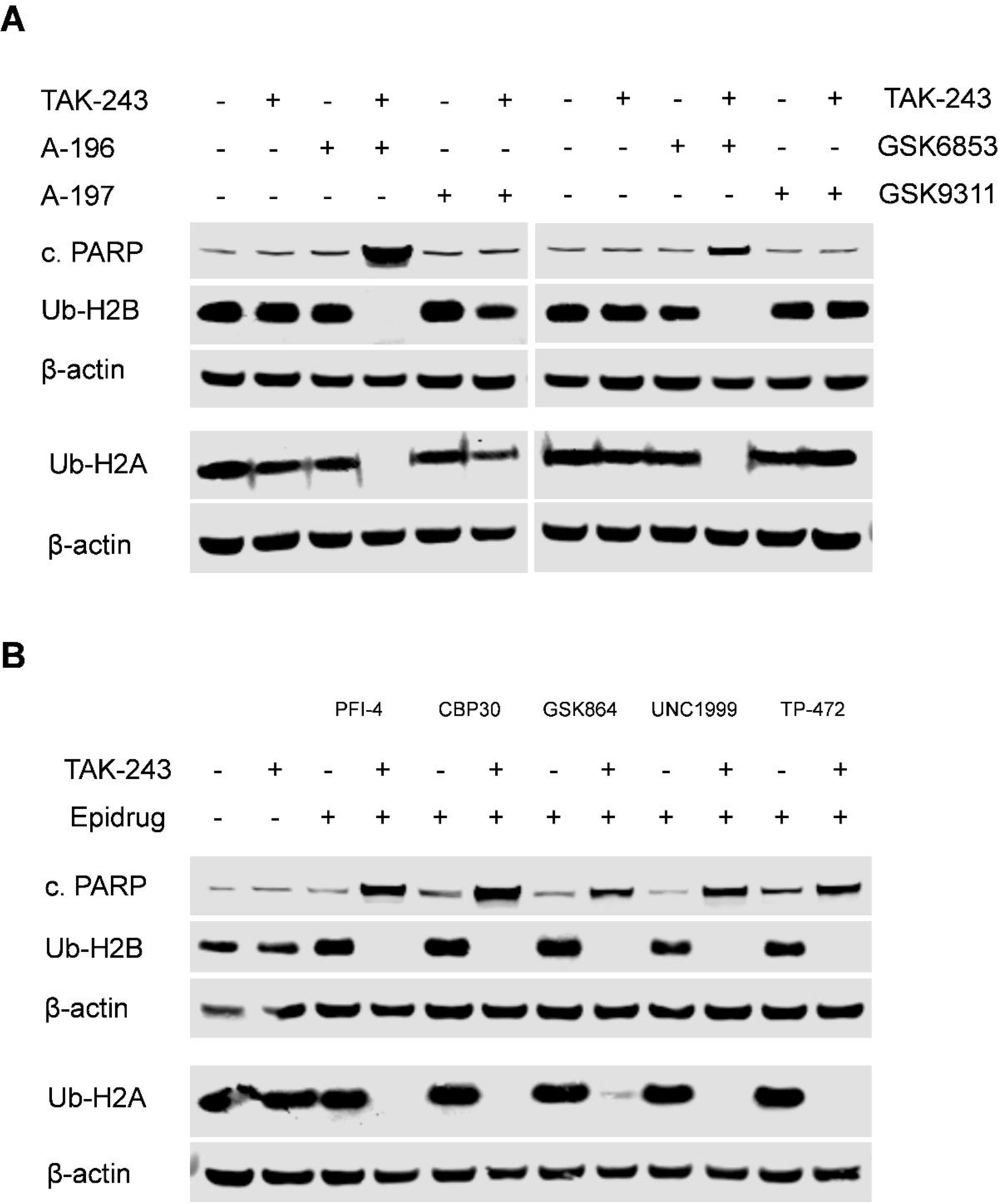
Potentiation of TAK-243 cytotoxicity is associated with reduced ubiquitylation and induction of apoptosis. RPMI 8226 cells were pre-treated with DMSO or epigenetic probes A-196, GSK6853 (or their negative controls A-197 and GSK9311, respectively) (A), PFI-4, SGC-CBP30, GSK-864, UNC1999, and TP-472 (B) for 24h. Subsequently cells were treated with DMSO or TAK-243 (250 nM) for 8h. After treatment, whole cell lysates were prepared and levels of cleaved PARP (c. PARP), ubiquitylated H2A (Ub-H2A), and Ub-H2B were measured by immunoblotting. β-actin was used as a loading control.

### Potentiation induced by the screen hits is relatively selective for TAK-243

As per the combinatorial screen data, profound potentiation is not observed with other anticancer agents, suggesting potentiation is TAK-243-specific (**Fig. 1A and S1)**. To further confirm this observation, we tested cross-potentiation by treating RPMI 8226 myeloma cells with increasing concentrations of 12 mechanistically and structurally distinct anticancer agents alone and in combination with A-196 and subsequently measured viability by the Resazurin assay after 72h. In line with the screen data, no cross-potentiation was observed with 10 anticancer drugs, with only partial cross-potentiation with pevonedistat (2-fold reduction in the IC_50_) and mitoxantrone (5-fold reduction in the IC_50_) (**Fig. S4**). Of note, pevonedistat is structurally and mechanistically related to TAK-243 as it is an adenosine sulfamate that selectively inhibits the NEDD8-activating enzyme (NAE) involved in regulating the ubiquitylation of a subset of cellular proteins^30^. Taken together, these data confirm the screen findings and suggest potentiation is relatively selective for TAK-243 particularly in terms of the magnitude of potentiation.

### Potentiation of TAK-243 cytotoxicity is associated with increased TAK-243 binding to UBA1

TAK-243 inhibits UBA1 resulting in the abrogation of ubiquitylation and induction of cell death^22, 29^. To gain mechanistic insights into the mechanism of chemical probe-induced potentiation of TAK-243 cytotoxicity, we treated RMPI 8226 cells with compounds identified from the screen for 24h and assessed the levels of UBA1, ubiquitin, Ub-H2A and global ubiquitylated proteins by immunoblotting. Treatment with different potentiators did not induce significant changes in the abundance of these proteins, suggesting there are no molecular changes in the ubiquitin system that may account for the observed potentiation (**Fig. S5**).

TAK-243 is a substrate of breast cancer resistance protein (BCRP/ABCG2), a member of the ATP-binding cassette (ABC) efflux transporters commonly involved in multi-drug resistance (MDR) by active transport of drugs out of the cells^31, 34^. Thus, we hypothesized downregulation of ABCG2 secondary to epigenetic changes caused by these probes may be responsible for the observed potentiation. However, we observed no changes in ABCG2 levels suggesting epigenetic changes elicited by these chemical probes had no impact on ABCG2 expression (**Fig. S5**).

Alternatively, these chemical probes may directly inhibit ABCG2 resulting in reduced efflux and increased intracellular TAK-243 accumulation. To test this hypothesis, we used the cellular thermal shift assay (CETSA) to assess the extent of TAK-243 binding to its target UBA1. CETSA is a cell-based biophysical assay used to evaluate the thermostabilization of proteins after binding to chemical ligands. Thus, higher CETSA signals correspond to higher thermostability reflecting an increased binding of the ligand to its target^21^. To perform CETSA, we pre-treated RPMI 8226 cells with A-196 for 48h followed by treatment with increasing concentrations of TAK-243 for 1h and subsequent evaluation of thermal shift of UBA1 using immunoblotting. Consistent with data from the cytotoxicity assays, A-196 treatment was associated with significant elevations in TAK-243 binding to UBA1 up to 500 nM, as evidenced by higher UBA1 signal in the soluble fraction measured by immunoblotting, reflecting increased intracellular drug accumulation (**Fig. 4A**). Similarly, treatments with SGC-CBP30, PFI-4, GSK-864, and GSK-6853 for 48h were associated with significant elevations in TAK-243 binding to UBA1 (**Fig. 4B**). Conversely, treatments with negative controls of A-196 (A197 and SGC-2043) and GSK-6853 (GSK-9311) were not associated with significant changes in TAK-243 binding (**Fig. 4B**). Similar results were observed after pre-treatment with chemical probes for only 4h, suggesting direct inhibitory activity (**Fig. 4B**). Of note, the mRNA expression of ABCG2 displayed a statistically significant positive correlation with the magnitude of TAK-243 potentiation observed with A-196 (**Fig. 2 and 4C**). These data suggest the observed potentiation is not attributable to on-target epigenetic effects of the chemical probes. To confirm this hypothesis, we used distinct chemical probes (also known as orthogonal probes) that inhibit the same epigenetic targets (**Fig. S6**). Specifically, we treated RPMI 8226 cells with increasing concentrations of TAK-243 alone and in combination with these orthogonal probes and assessed viability by the Resazurin assay. All probes, except for GSK5959, did not result in significant reductions in the IC_50_ of TAK-243 **(Fig. S6B-F)**. Of note, GSK5959 shared significant structural similarity with PFI-4 and GSK6853, perhaps explaining the retention of the off-target effects on ABCG2 **(Fig. S6B)**. Taken together, our data suggest these epigenetic probes may exert their potentiation of TAK-243 cytotoxicity through an off-target mechanism by ABCG2 inhibition rather than inducing changes in ABCG2 expression via an epigenetic mechanism.

**Figure 4.**
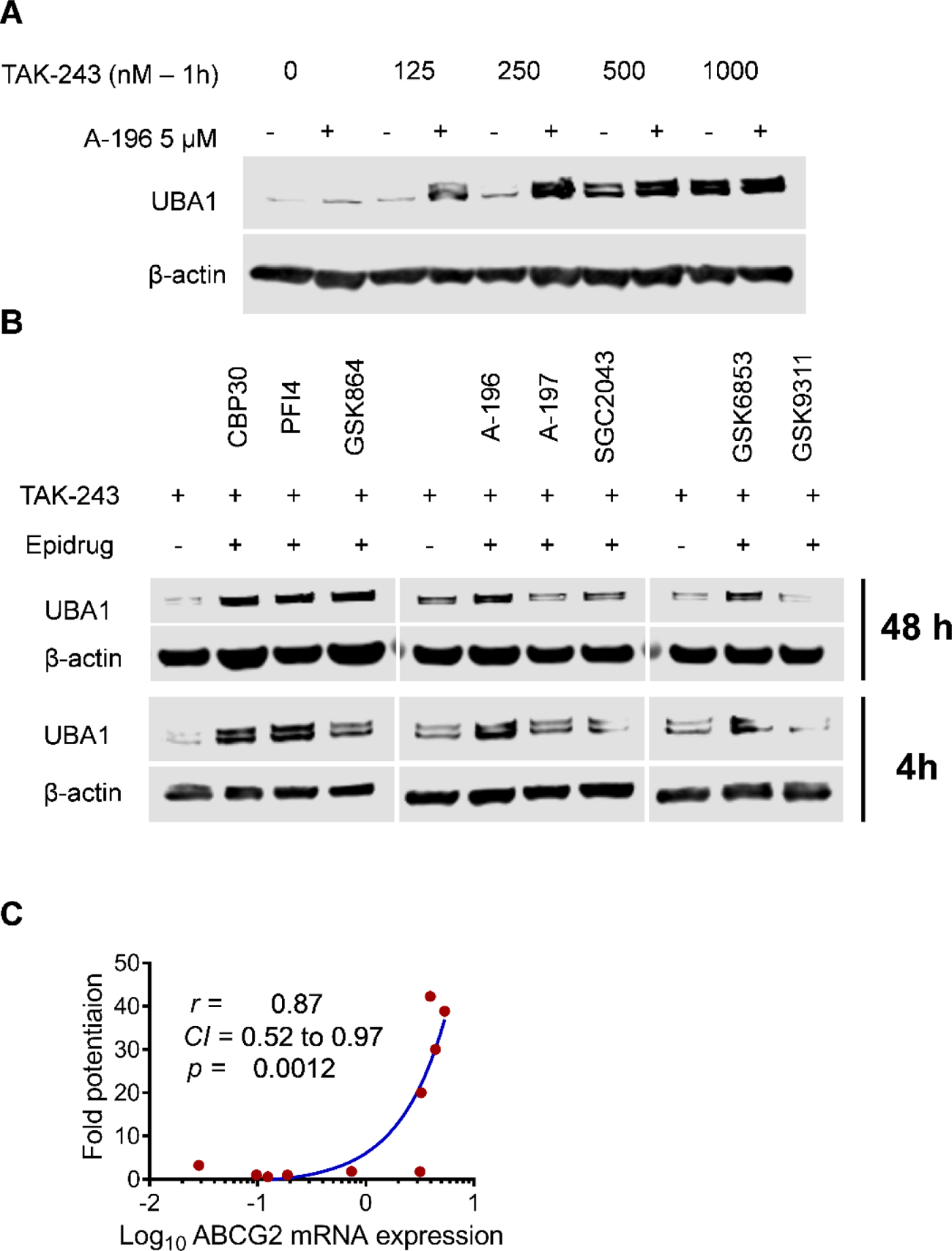
Epigenetic probes, but not their negative controls, increase TAK-243 binding to its target UBA1. A) RPMI 8226 cells were pretreated with A-196 (5 µM) for 48h followed by treatment with increasing concentrations of TAK-243 (125-1000 nM) for 1h. The cellular thermal shift assay (CETSA) was conducted in which cells were washed, resuspended in PBS, and heated for 3 min at 54°C followed by lysis through 3 freeze-thaw cycles. Supernatants were collected and UBA1 levels were assessed by immunoblotting. B) The CETSA assay was conducted after treatment with TAK-243 alone and in combination with chemical probes and their negative controls after pre-treatment with chemical probes for 48h (upper panel) or 4h (lower panel). β-actin was used as a loading control. C) Correlation of mRNA expression of ABCG2 in 10 cell lines as obtained from the DepMap database (https://depmap.org/portal/) and the fold-potentiation of TAK-243 cytotoxicity observed with A-196. *r* denotes correlation coefficient and *CI* denotes confidence interval.

### ABCG2 knockdown sensitizes cells to TAK-243 and abrogates potentiation observed with different chemical probes

To assess the functional importance of ABCG2 in chemical probe-induced potentiation of TAK-243 cytotoxicity, we knocked down *ABCG2* in A549 and RPMI 8226 cells and confirmed knockdown by immunoblotting (**Fig. 5A**). We subsequently treated control and knockdown cells with increasing concentrations of TAK-243 alone and in combination with different epigenetic probes. In control cells, as previously observed, combinatorial treatment with different screen hits resulted in profound reductions in the IC_50_ of TAK-243 by up to 14- and 40-fold in A549 and RPMI 8226 cells, respectively (**Fig. 5B-C**). In contrast, *ABCG2* knockdown significantly abrogated the potentiation as evidenced by much lower reductions in the IC_50_ values (**Fig. 5D-I**). Taken together, these data confirm ABCG2 as an efflux transporter for TAK-243 and implicate ABCG2 as a candidate target that is inhibited by the screen hits resulting in potentiation of TAK-243 cytotoxicity.

**Figure 5.**
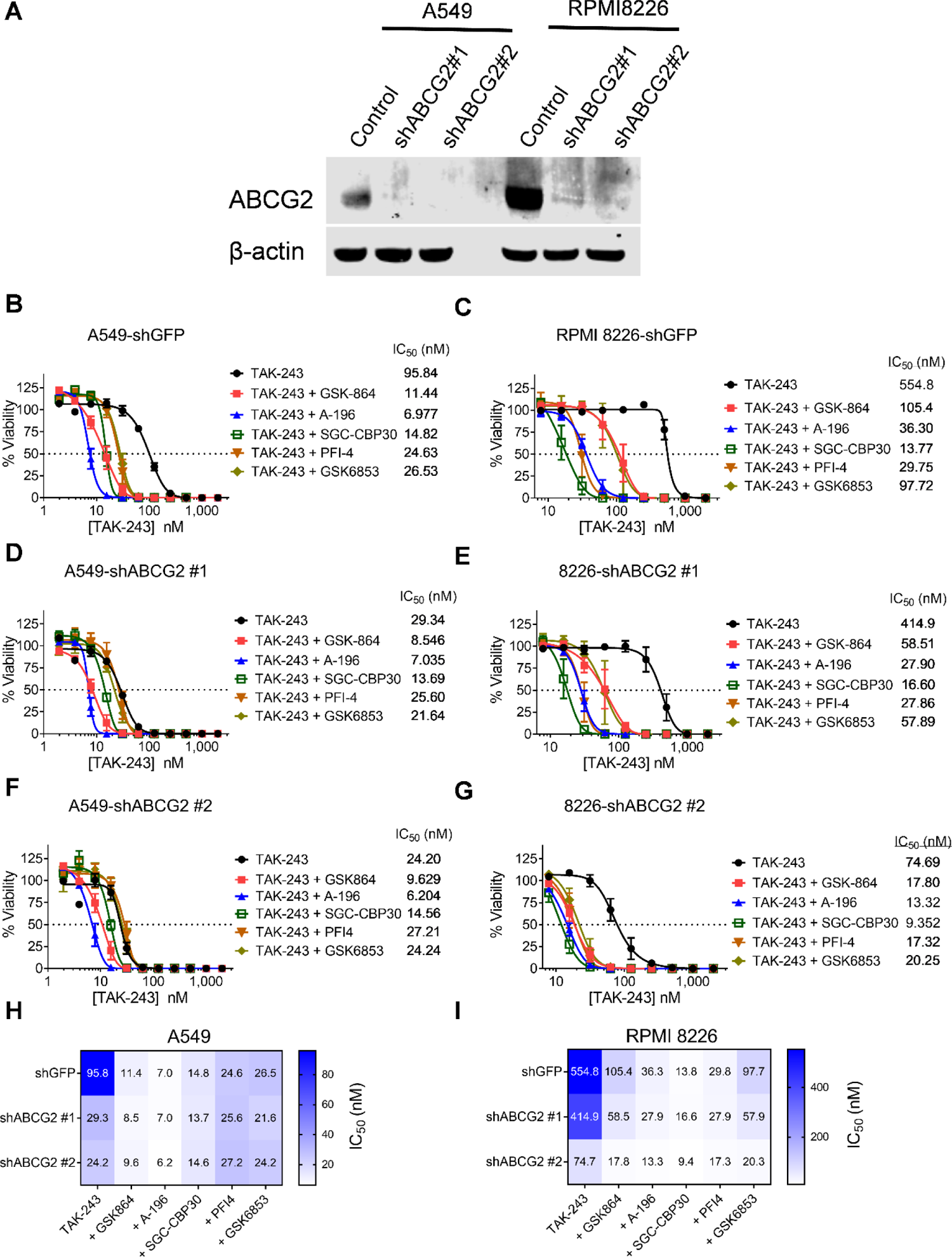
*ABCG2* knockdown sensitizes cells to TAK-243 and abrogates potentiation observed with different chemical probes. A) A549 and RPMI-8226 cells were transduced with non-targeting and 2 distinct shRNAs targeting *ABCG2* and cell lysates were collected followed by measuring levels of ABCG2 by immunoblotting. B-H) Control and *ABCG2*-kncokdown A549 (B, D, F) and RPMI 8226 (C, E, G) cells were treated with increasing concentrations of TAK-243 alone and in combination with different chemical probes, and viability was measured by the Resazurin assay after 72h. Insert: median inhibitory concentration (IC_50_) values. All data points represent the mean ± SEM of at least 3 independent experiments H, I) Heatmaps depicting the IC_50_ values of TAK-243 alone and in combination with different chemical probes in A549 (H) and RPMI 8226 (I) cell lines.

### Chemical probes share significant structural similarity with TAK-243 and independently potentiate the cytotoxicity of ABCG2 substrates

To explain their the ABCG2-inhibitory activities, we analyzed the structural features shared among the identified chemical probes and TAK-243 known to serve as an ABCG2 substrate^31, 35^. As previously reported, TAK-243 is an adenosine sulfamate that comprises three main structural moieties: 1) cyclopentane moiety; 2) purine nucleus; and 3) aromatic/heterocyclic extension **(Fig S7A)**^30^. Similarly, the chemical probes possess chemical moieties that are closely related or analogous to those comprised by TAK-243, explaining their potential ABCG2-inhibitory activities **(Fig S7A)**. To verify these inhibitory activities independent of TAK-243, we treated RPMI 8226 cells with increasing concentrations with the ABCG2 substrates, mitoxantrone and pevonedistat, alone and in combination with different epigenetic probes. Although not as profound as observed with TAK-243, the chemical probes potentiated the cytotoxicity of mitoxantrone (2- to 5-fold) and pevonedistat (2-fold), confirming the ABCG2-inhibitory activity of these chemical probes **(Fig. S7B-C)**. The profound potentiation observed with TAK-243 may stem from the relative differences in affinities to ABCG2, the pharmacodynamics of cytotoxicity and the presence of other transporters for which mitoxantrone/pevonedistat is a substrate. Taken together, these data show the significant structural similarities between TAK-243 and chemical probes and consolidate their potential ABCG2-inhibitory activities.

### Chemical probes showed a comparable docking into the ABCG2 substrate-binding site as known ABCG2 inhibitors

To further investigate the mechanism of action of probes that potentiate TAK-243 cytotoxicity, we used computational docking to test whether the compound could fit in the ligand binding site of ABCG2. Docking was conducted on both open and closed conformations of the protein. In the inward-open conformation (PDB: 6ETI, 7NEQ)^23, 24, 36–38^, the transmembrane domains of two ABCG2 monomers form a substrate-binding cavity open to the cytoplasm, whereas the turnover-closed conformation (PDB: 7OJ8)^25^, in which the endogenous steroid estrone sulfate (E1S) is bound, is closed to the cytoplasm.

Chemical probes were docked to ABCG2 with Glide (Schrodinger). Docking scores ranged from −7 to −10 kcal/mol across the ABCG2 structures, similar to scores of known inhibitors and substrates such as MZ29^23^, tariquidar (EC_50_= 0.08 ± 0.01 µM)^24^, and E1S (EC_50_= 15.7 ± 0.9 µM)^36^, suggesting the probes may fit in the ABCG2 substrate-binding site in both the inward-open and turnover-closed conformations (**Table S4, Fig. 6A-C**). Additionally, docking analysis predicted that the polycyclic core of TAK-243 and the probes share key stacking interactions with Phe439 in the ABCG2 binding pocket (**Fig. 6D**).

**Figure 6.**
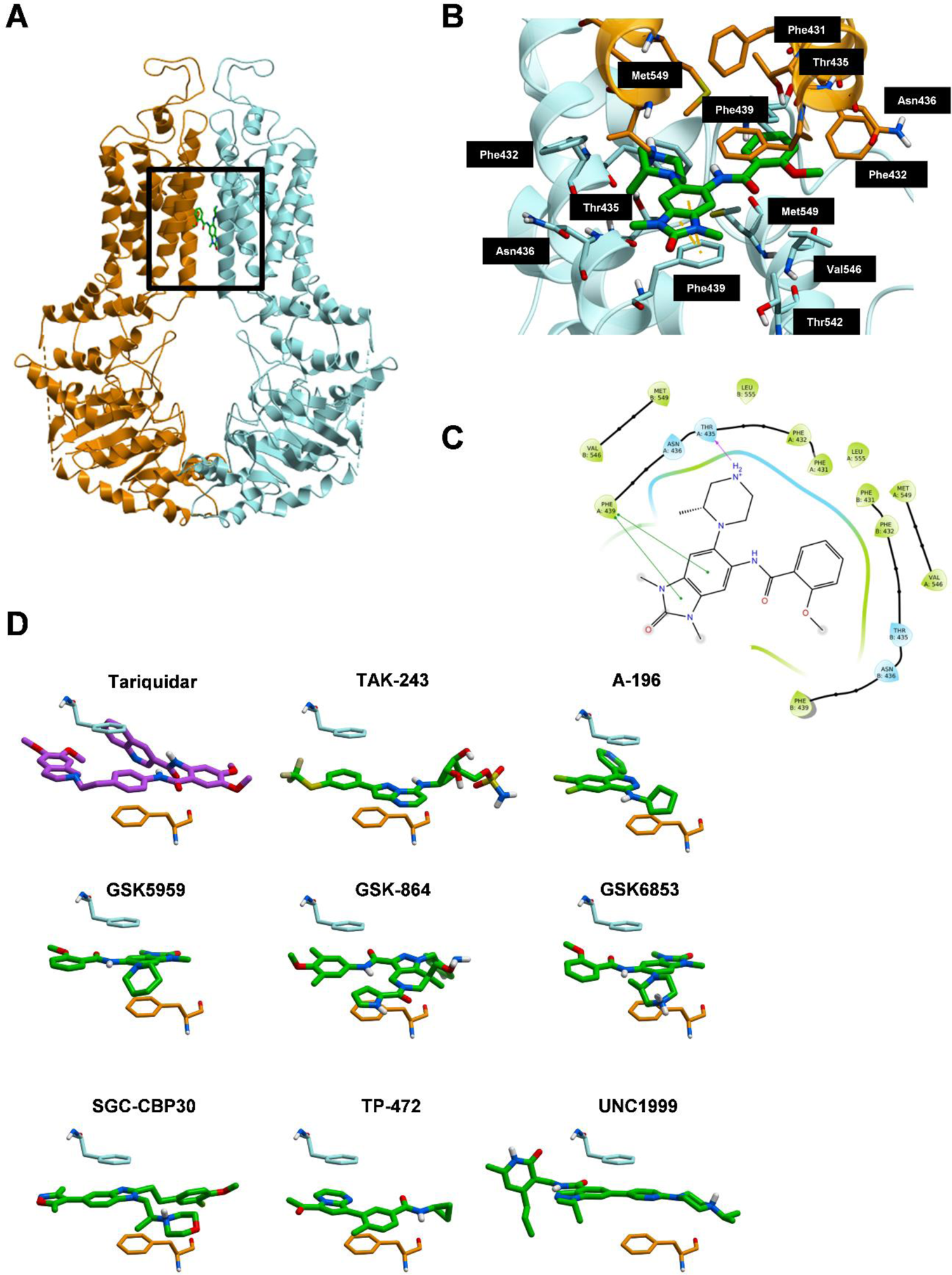
Docking analysis of chemical probe interaction with ABCG2. (A-C) Docking model of GSK6853 bound to ABCG2 (A) Overview of best-scoring pose of GSK6853 in the binding pocket of ABCG2 protein (PDB: 6ETI). ABCG2 is displayed as a ribbon representation (chain A: blue, chain B: orange). GSK6853 is displayed in sticks (green). (B) Details of interaction between ABCG2 and GSK6853. Select residues are displayed as sticks (PDB: 6ETI). Residues in chain A and B are labeled on black and grey backgrounds, respectively. Hydrogen bonds are displayed as purple dashed lines and π-π stacking interactions are displayed as green dashed lines. (C) 2D ligand interaction map between GSK6853 and ABCG2 binding pocket generated with Maestro (Schrodinger). Amino acids in the binding pocket are colored bubbles (green: hydrophobic; blue: polar). Purple solid lines with arrows indicate hydrogen bonds. Green solid lines indicate π-π stacking. D) Docked poses of TAK-243 and probes share interactions with ABCG2 (PDB: 7NEQ). Phe439 (chain A: blue, chain B: orange), crystallized ligand (purple) and docked ligands (green) are displayed as sticks.

Negative controls for each probe were also docked and generated less favorable scores in the case of GSK9311 (negative control of GSK6853) and UNC2400 (negative control of UNC1999) but as favorable in other cases. Considering the relatively low resolution of available ABCG2 electron microscopy structures, we believe that estimating the affinity of these ligands computationally would be an unreliable exercise. Nevertheless, these data suggest that ABCG2 could indeed accommodate, and potentially be inhibited by the chemical probes tested in this work.

### Potentiation of TAK-243 cytotoxicity can serve as a cell-based assay of ABCG2-inhibitory activity

Our screen has identified TAK-243 as an anticancer agent whose cytotoxicity is profoundly potentiated by off-target ABCG2-inhibitory activities of epigenetic probes. In contrast to other ABCG2 substrates such as mitoxantrone and pevonedistat, TAK-243 induced rapid and profound cytotoxicity upon inhibition of ABCG2. Thus, TAK-243 along with ABCG2-overexpressing cancer cell lines such as A549 and RPMI 8226 can be potentially used to robustly assay the ABCG2-inhibitory activity of novel compounds. To test this hypothesis, we treated RPMI 8226 cells with increasing concentrations of epigenetic probes alone and in combination with TAK-243. For TAK-243, we selected a concentration of 250 nM that does not affect the viability of RPMI 8226 when used alone but would certainly abolish the viability of these cells upon complete ABCG2 inhibition. As assessed by the Resazurin assay after 72h of incubation, all epigenetic probes did not significantly affect the viability when used alone except for TP-472 that showed significant cytotoxicity at 10 µM. When combined with TAK-243, the epigenetic probes reduced viability in a concentration-dependent manner, reflecting the relative potencies of their ABCG2-inhibitory activities rather than their cytotoxicity (**Fig. 7A-G**). In parallel, we similarly treated cells with the ABCG2 inhibitor Ko143 and the P-glycoprotein inhibitor zosuquidar to serve as positive and negative controls, respectively (**Fig. 7H-I**)^39–41^. As a surrogate measure of these potencies, we calculated the apparent IC_50_ values of combinatorial treatments and ranked the epigenetic inhibitors in order of these values. As expected, Ko143 displayed the highest potency with an IC_50_ value of approximately 50 nM, with no significant changes observed with Zosuquidar (**Fig. 7J**). Of the epigenetic inhibitors, PFI-4, A-196, SGC-CBP30, and UNC1999 displayed sub-micromolar IC_50_ values from 250 to 850 nM, while other inhibitors displayed micromolar IC_50_ values from 1.5-3 µM (**Fig. 7J**). Taken together, these data validate the use of TAK-243 as a tool that can be used in cell-based assays to measure the potential ABCG2-inhibitory activities of novel compounds in a quantitative manner.

**Figure 7.**
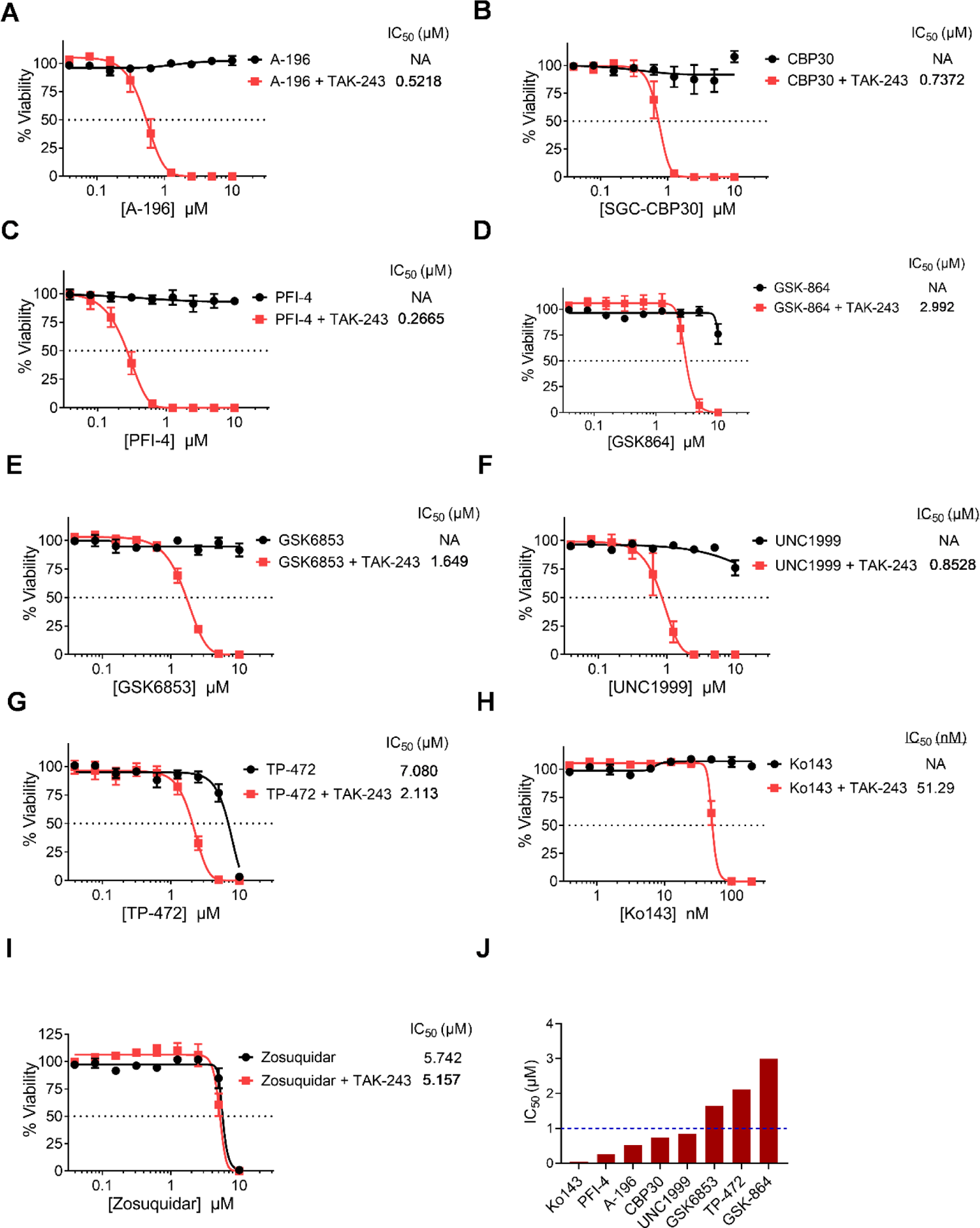
Potentiation of TAK-243 cytotoxicity can serve as a cell-based assay of ABCG2 inhibition. A-I) Concentration-response curves of epigenetic probes (A-G), the ABCG2 inhibitor Ko143 (H), and the P-gp inhibitor Zosuquidar (I) after treatment of RPMI 8226 cells alone and in combination with TAK-243 (250 nM) for 72h. Insert: median inhibitory concentration (IC_50_) values. J) Bar chart ranking the apparent IC_50_ values of epigenetic probes and Ko143 after combination with TAK-243 (250 nM) in RPMI 8226 cells.

## Discussion

Epigenetic probes represent a valuable chemical biology tool that can be used in combinatorial screens to explore synthetic lethal interactions with anticancer agents^42, 43^. Such screens may provide further insights into the regulation of anticancer response and potentially identify effective anticancer drug combinations^43^. In this study, we exploited the SGC toolbox of epigenetic probes and a panel of anticancer agents to conduct a combinatorial screen. Using this approach, we identified several epigenetic probes that potentiate the cytotoxicity of TAK-343.

TAK-243 is a first-in-class UBA1 inhibitor that displays preclinical efficacy in a broad spectrum of solid and hematologic malignancies including colorectal cancer^29^, glioblastoma^44^, renal cell carcinoma^45^, small-cell lung cancer^46^, acute myeloid leukemia^22^, multiple myeloma^47^, B-cell lymphoma^48^, and chronic lymphocytic leukemia^49^. Additionally, it has been advanced to phase 1 clinical trials in advanced malignancies and acute myeloid leukemia^30^.

Our study demonstrated the potentiation of TAK-243 cytotoxicity was associated with profound reduction in histone ubiquitylation and induction of PARP cleavage, consistent with the molecular effects reported with TAK-243 in previous studies^22, 29^.

Among anticancer agents tested, profound potentiation was only observed with TAK-243 by approximately 20% of the epigenetic probes used in the screen. Additionally, the potentiation was not cancer specific as it was observed in cell lines of different origin. This high hit rate across functionally diverse epigenetic regulators, along with the lack of cancer specificity suggested an off-target mechanism may be responsible for the observed potentiation.

Negative controls are important chemical biology tools that are used along with chemical probes to exclude potential off-target effects^32^. With their closely related structures, these negative controls lack on-target effects and should ideally retain off-target effects of the chemical probes^32^. Our cytotoxicity and molecular experiments showed that most negative control compounds used in the study did not phenocopy the off-target effects observed with epigenetic probes. While seemingly contradictory to our hypothesis, a recent computational study showed that chemical probe negative controls may lose off-target as well as on-target activities, compromising their utility in excluding off-target effects^50^. Nonetheless, structurally distinct orthogonal inhibitors of the targeted epigenetic regulators did not elicit significant TAK-243 potentiation; thus, indicating the importance of such orthogonal compounds in functional studies.

Our study showed no significant changes in the expression levels of molecules involved in ubiquitylation including UBA1 and ubiquitin, nor in the abundance of ubiquitylated proteins, supporting an off-target potentiation mechanism. Several molecular biomarkers have been reported to determine the sensitivity of TAK-243 in different contexts including ABCG2, SLFN11, GRP78 and YAP1^31, 44, 46, 51^. Herein, we demonstrated that the identified chemical probes induced rapid increases in intracellular TAK-243 concentrations and potentiated the cytotoxicity of the ABCG2 substrates pevonedistat and mitoxantrone with no changes in ABCG2 expression levels. Thus, these observations implicated ABCG2 as a target that is directly inhibited by these chemical probes, which we confirmed through genetic knockdown and docking experiments.

ABCG2 is an ATP-dependent ABC efflux transporter reported to play an important role in the resistance to chemotherapeutic agents including TAK-243^31, 35^, pevonedistat^46^, mitoxantrone^52, 53^, and doxorubicin^53^. Due to the high potency of TAK-243 and essential role of ubiquitylation in almost all cell biological functions, the doubling or tripling of intracellular TAK-243 concentrations by these chemical probes is anticipated to result in rapid and profound reductions in viability, as opposed to mitoxantrone or pevonedistat that have different cytotoxicity pharmacodynamics.

Docking analysis indicate that the probes can potentially fit in the hydrophobic cavity located at the center of the ABCG2 transmembrane domain. The epigenetic probes have structural similarities with TAK-243, an ABCG2 substrate, including a polycyclic core and an aromatic extension. It has been shown that depending on the size of the compounds, one or two inhibitor molecules can bind in the ABCG2 pocket^23^. The docking study considered binding of one ligand to ABCG2 but does not discount the fact that some ligands may bind with a different stoichiometry. Recently, ABCG2 has been reported to mediate the resistance to GSK1070916, an investigational anticancer agent that targets Aurora kinase^54^. By docking analysis, the authors demonstrated the interaction between GSK1070916 and the substrate-binding site of ABCG2. Interestingly, GSK1070916 shares similar structural features with TAK-243 and the epigenetic probes identified in our study. Accordingly, we speculate anticancer agents with these structural features may serve as ABCG2 substrates and thus testing for such activity needs to be prioritized during the preclinical development of these drug classes.

In our study, we proposed and validated the use of TAK-243 as a tool to quantify ABCG2-inhibitory activities of novel compounds in cell-based assays. The drugs commonly used in resistance assays, such as doxorubicin and mitoxantrone, do not exhibit quantitative reductions of the viability upon ABCG2 inhibition^34^. In contrast, TAK-243 displays a more robust cytotoxicity that correlates with the extent of ABCG2 inhibition. In our validation experiments, the IC_50_ values of TAK-243 accurately reflected the ABCG2-inhibitory activities of the epigenetic probes. Thus, potentiation of TAK-243 can potentially serve as indirect quantitative assay of the potency of ABCG2 inhibition by using appropriate positive controls such as Ko143.

In summary, our study identifies epigenetic probes that profoundly potentiate TAK-243 cytotoxicity through off-target ABCG2 inhibition that is not excluded by negative chemical controls but can be addressed by using structurally distinct orthogonal probes. These findings may also suggest common structural features for probes with potential inhibitory activity of ABCG2, an important MDR transporter for many drugs from different therapeutic classes Moreover, we propose the use of TAK-243 in a cell-based assay that can reliably quantify ABCG2 inhibition by drug candidates.

## Author contributions

S.H.B. and Y.Y. conducted experiments and analyzed data; M.K.M. performed docking analysis; M.S. supervised docking analysis; A.D.S, C.H.A. and D.B.L supervised research, data analysis and interpretation; S.H.B. and D.B.L conceived the project and designed the experiments; S.H.B. wrote the first draft of the manuscript. All authors reviewed and edited the manuscript.

## Acknowledgments

S.H.B is a Mitacs Elevate Fellow. The Structural Genomics Consortium is a registered charity (no: 1097737) that receives funds from Bayer AG, Boehringer Ingelheim, Bristol Myers Squibb, Genentech, Genome Canada through Ontario Genomics Institute [OGI-196], EU/EFPIA/OICR/McGill/KTH/Diamond Innovative Medicines Initiative 2 Joint Undertaking [EUbOPEN grant 875510], Janssen, Merck KGaA (aka EMD in Canada and US), Pfizer and Takeda.

## Supplementary Information

**Table S1.**
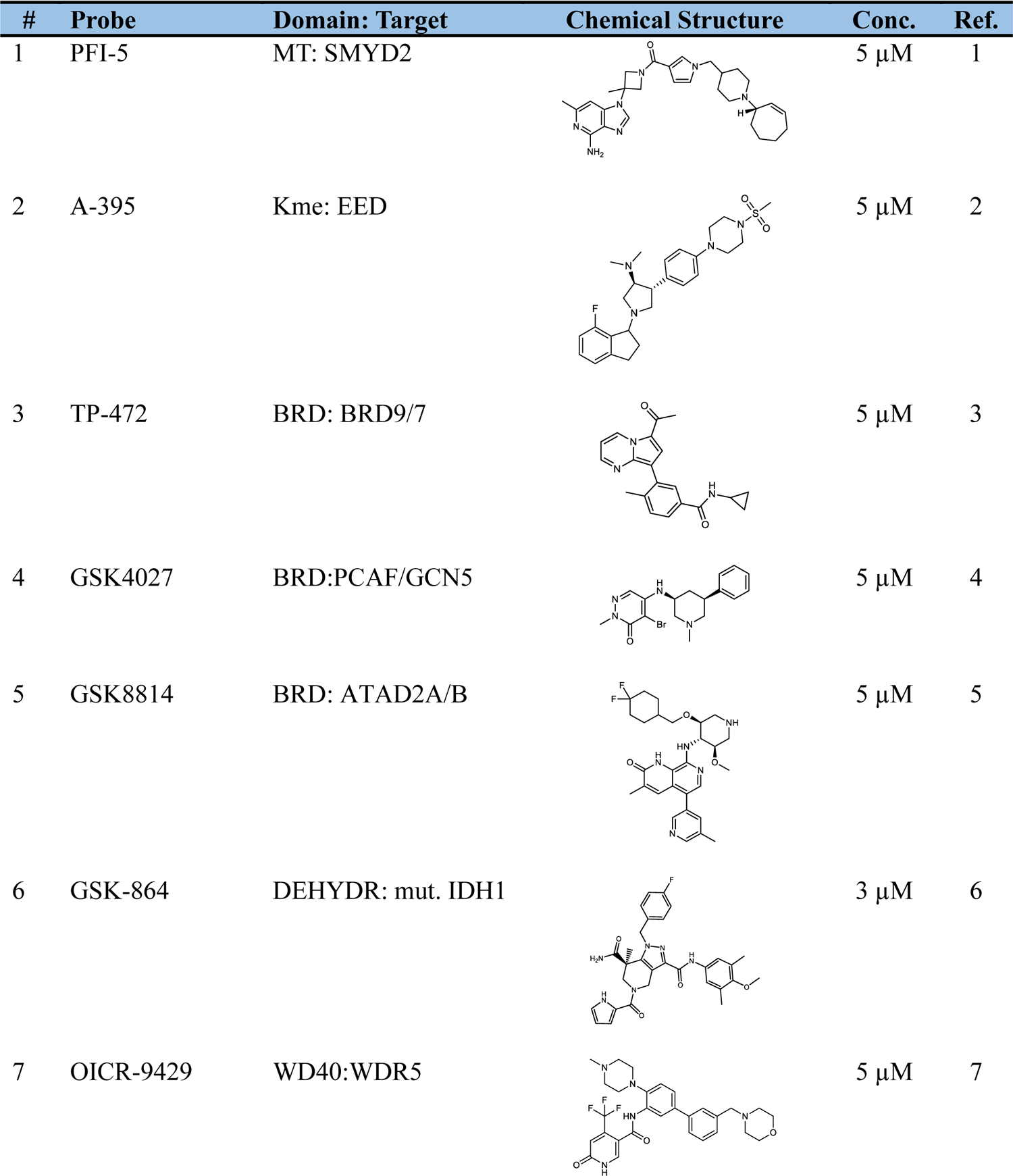
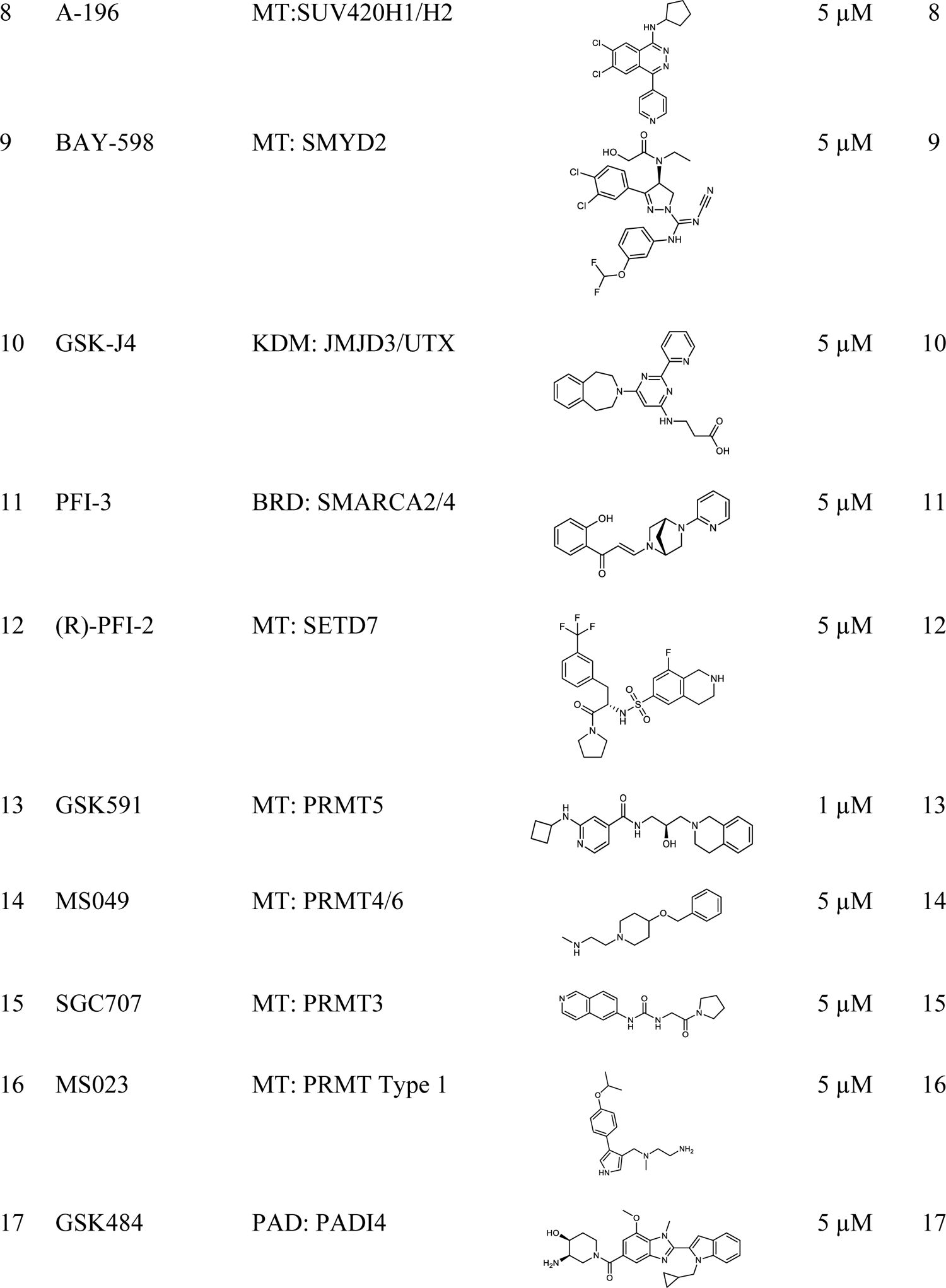
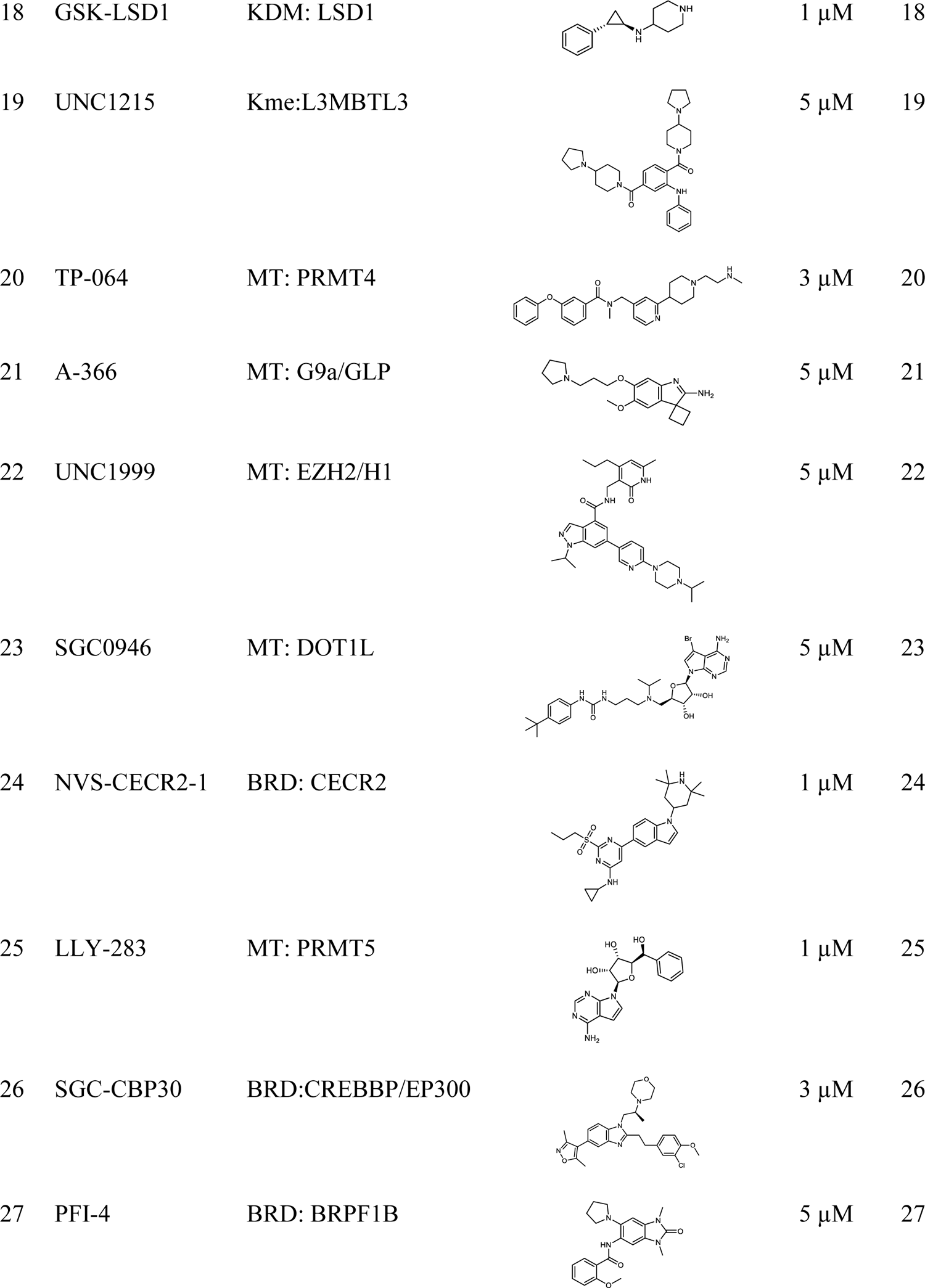
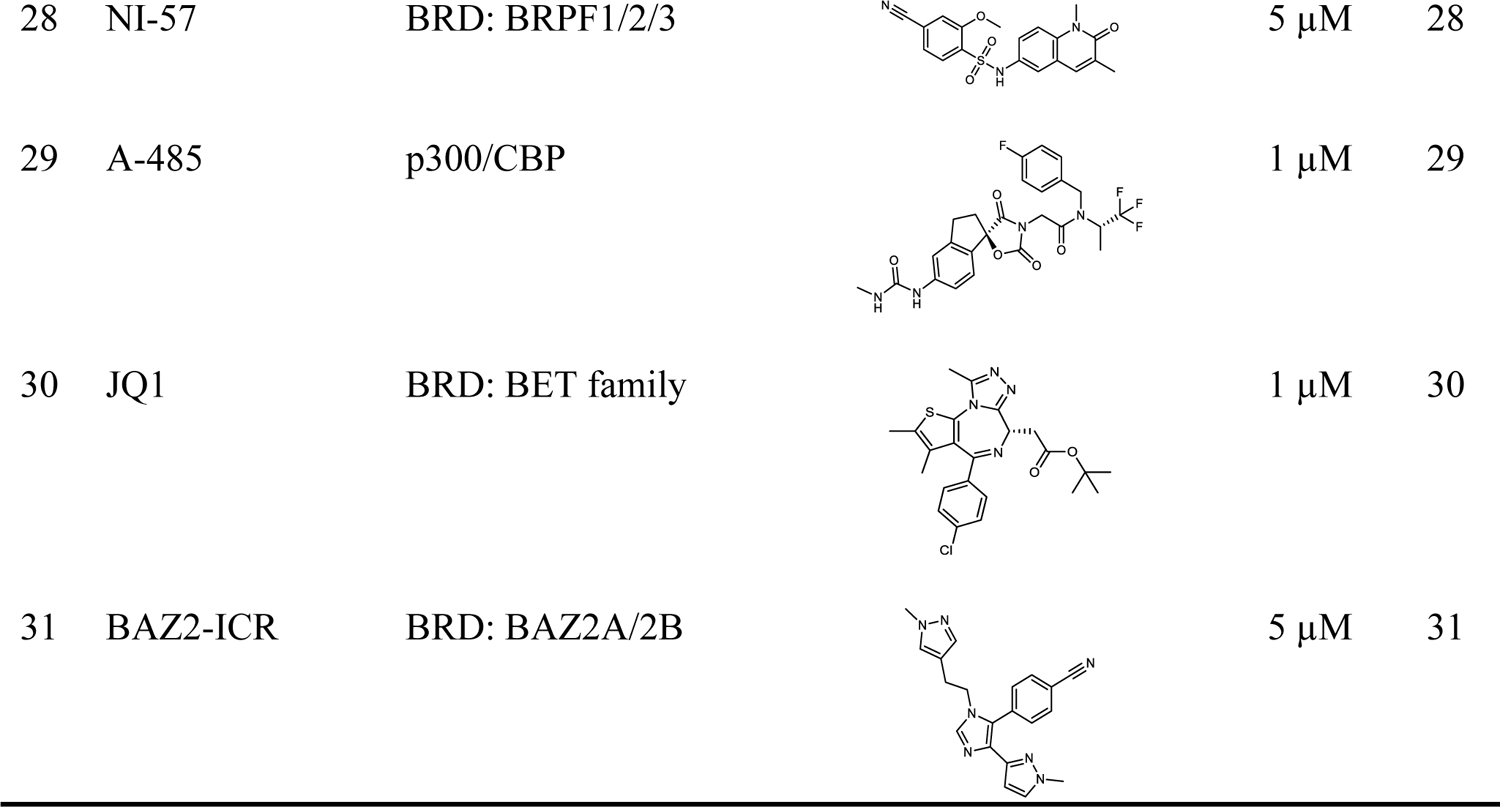
List of epigenetic probes used for the screen

**Table S2.**
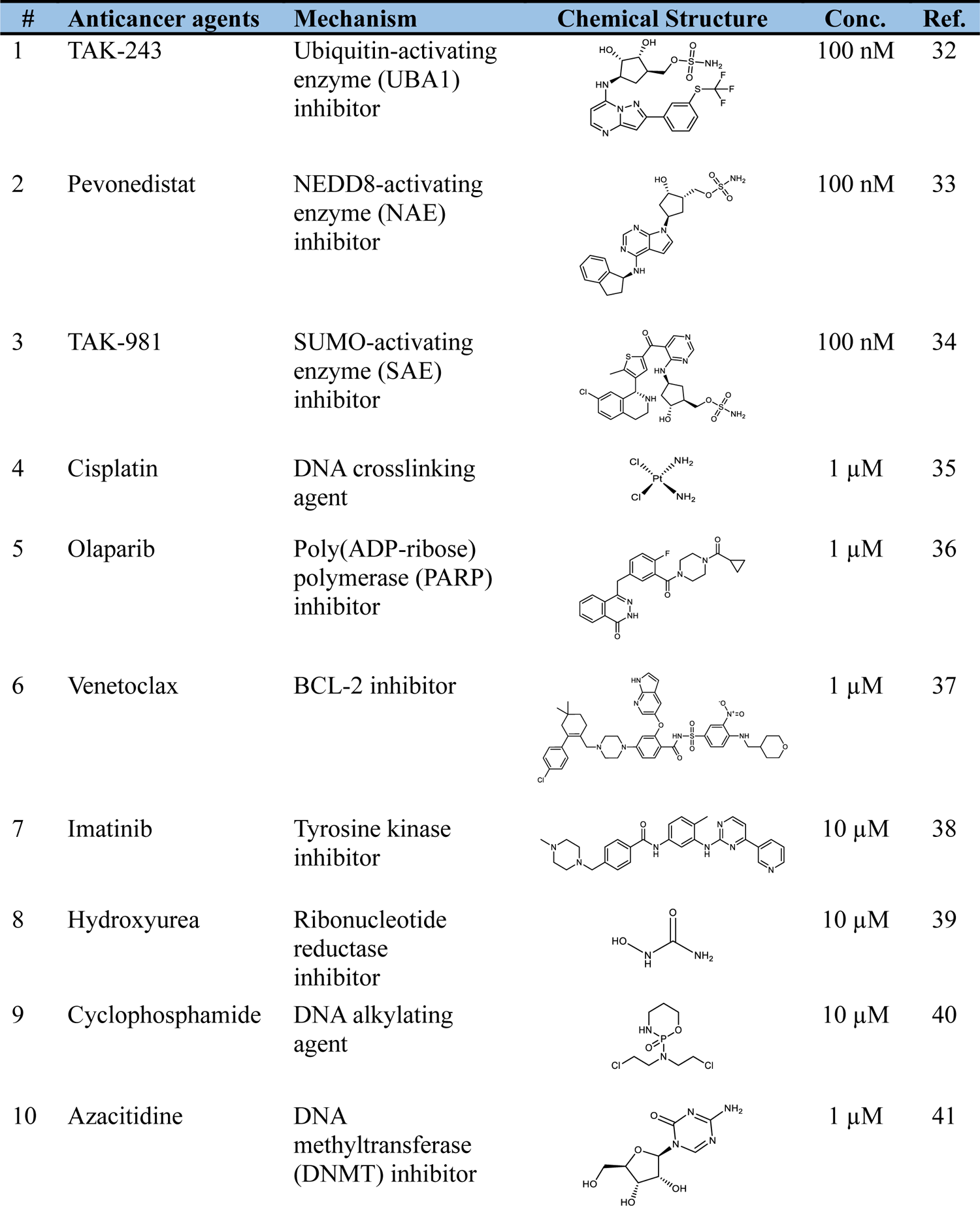
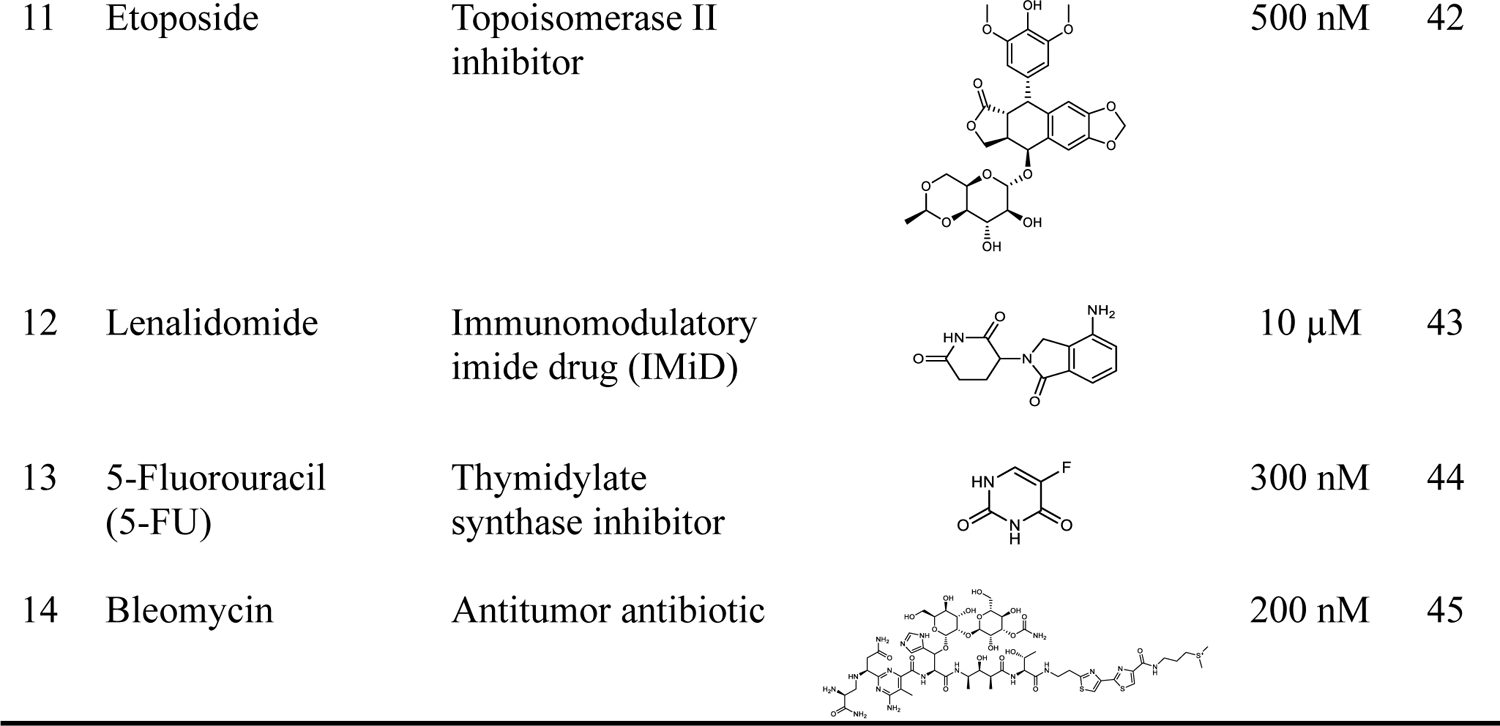
List of anticancer agents used for the screen

| # | Anticancer agents | Mechanism | Chemical Structure | Conc. | Ref. |
| --- | --- | --- | --- | --- | --- |
| 1 | TAK-243 | Ubiquitin-activating enzyme (UBA1) inhibitor |  | 100 nM | 32 |
| 2 | Pevonedistat | NEDD8-activating enzyme (NAE) inhibitor |  | 100 nM | 33 |
| 3 | TAK-981 | SUMO-activating enzyme (SAE) inhibitor |  | 100 nM | 34 |
| 4 | Cisplatin | DNA crosslinking agent | | 1 μM | 35 |
| 5 | Olaparib | Poly(ADP-ribose) polymerase (PARP) inhibitor | | 1 μM | 36 |
| 6 | Venetoclax | BCL-2 inhibitor | | 1 μM | 37 |
| 7 | Imatinib | Tyrosine kinase inhibitor | | 10 μM | 38 |
| 8 | Hydroxyurea | Ribonucleotide reductase inhibitor | | 10 μM | 39 |
| 9 | Cyclophosphamide | DNA alkylating agent | | 10 μM | 40 |
| 10 | Azacitidine | DNA methyltransferase (DNMT) inhibitor | | 1 μM | 41 |
| 11 | Etoposide | Topoisomerase II inhibitor |  | 500 nM | 42 |
| 12 | Lenalidomide | Immunomodulatory imide drug (IMiD) | | 10 μM | 43 |
| 13 | 5-Fluorouracil (5-FU) | Thymidylate synthase inhibitor |  | 300 nM | 44 |
| 14 | Bleomycin | Antitumor antibiotic |  | 200 nM | 45 |

**Table S3.** List of primary and secondary antibodies

| **Antibody** | **Source** | **Clone** | **Catalog#** | **Dilution** |
| --- | --- | --- | --- | --- |
| **β-actin** | Santa Cruz | C4 | sc-47778 | Primary: 1:1000<br>Secondary: 1:5000 |
| **ABCG2** | Cell Signaling | D5V2K | 42078 | Primary: 1:1000<br>Secondary: 1:5000 |
| **Cleaved PARP**<br>**(Asp214)** | Cell Signaling | D64E10 | 5625 | Primary: 1:1000<br>Secondary: 1:5000 |
| **Global ubiquitylated**<br>**proteins** | Millipore<br>Sigma | FK2 | 4263 | Primary: 1:1000<br>Secondary: 1:5000 |
| **UBA1** | Santa Cruz | B-5 | sc-271821 | Primary: 1:500<br>Secondary: 1:5000 |
| **Ubiquityl-Histone**<br>**H2A (Lys119)** | Cell Signaling | D27C4 | 8240 | Primary: 1:500<br>Secondary: 1:5000 |
| **Ubiquityl-Histone**<br>**H2B (Lys120)** | Cell Signaling | D11 | 5546 | Primary: 1:1000<br>Secondary: 1:5000 |
| **Ubiquitin** | Abcam | - | ab7254 | Primary: 1:1000<br>Secondary: 1:5000 |

**Table S4.** **Docking Scores (obtained with Glide SP) of chemical probes and negative controls to the open conformation (PDB: 6ETI, 7NEQ) and the closed conformation (PDB: 7OJ8) of ABCG2**

| Ligand | Score (kcal/mol) |  |  |
| --- | --- | --- | --- |
|  | 6ETI | 7NEQ | 7OJ8 |
| **TAK243** | -10.305 | -10.891 | -11.157 |
| **MZ29** | -8.531 |  |  |
| **Tariquidar** |  | -10.65 |  |
| **E1S** |  |  | -9.263 |
| **TP-472** | -8.442 | -8.483 | -9.842 |
| **TP-472N** | -8.61 | -7.886 | -8.825 |
| **A-196** | -8.012 | -7.603 | -8.468 |
| **A-197** | -8.871 | -8.442 | -9.466 |
| **SGC-2043** | -6.802 | -7.395 | -7.654 |
| **UNC1999** | -10.31 | -9.743 | -10 |
| **UNC2400** | -9.641 | -8.162 | NC |
| **GSK6853** | -9.774 | -9.627 | -9.586 |
| **GSK9311** | -7.468 | -7.12 | -8.944 |
| **GSK-864** | -9.939 | -9.899 | -10.71 |
| **SGC-CBP30** | -7.104 | -7.75 | -9.706 |
| **PF-I4** | NC | NC | NC |
| **GSK5959** | -8.910 | -8.757 | -9.573 |

NC- docking produced no reasonable poses; Positive control: blue background; Probes: grey background; Negative controls: white background

**Figure S1.**
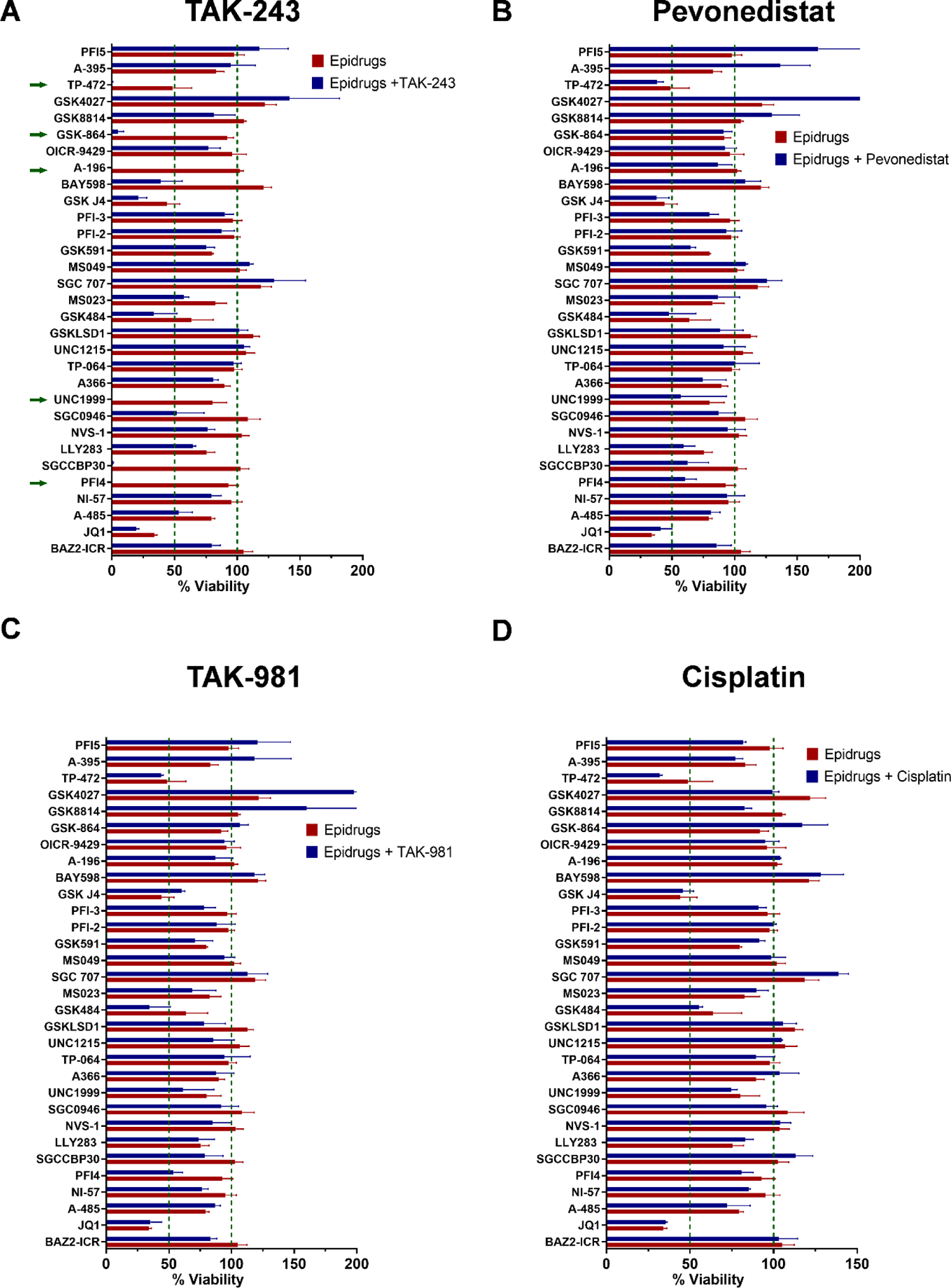
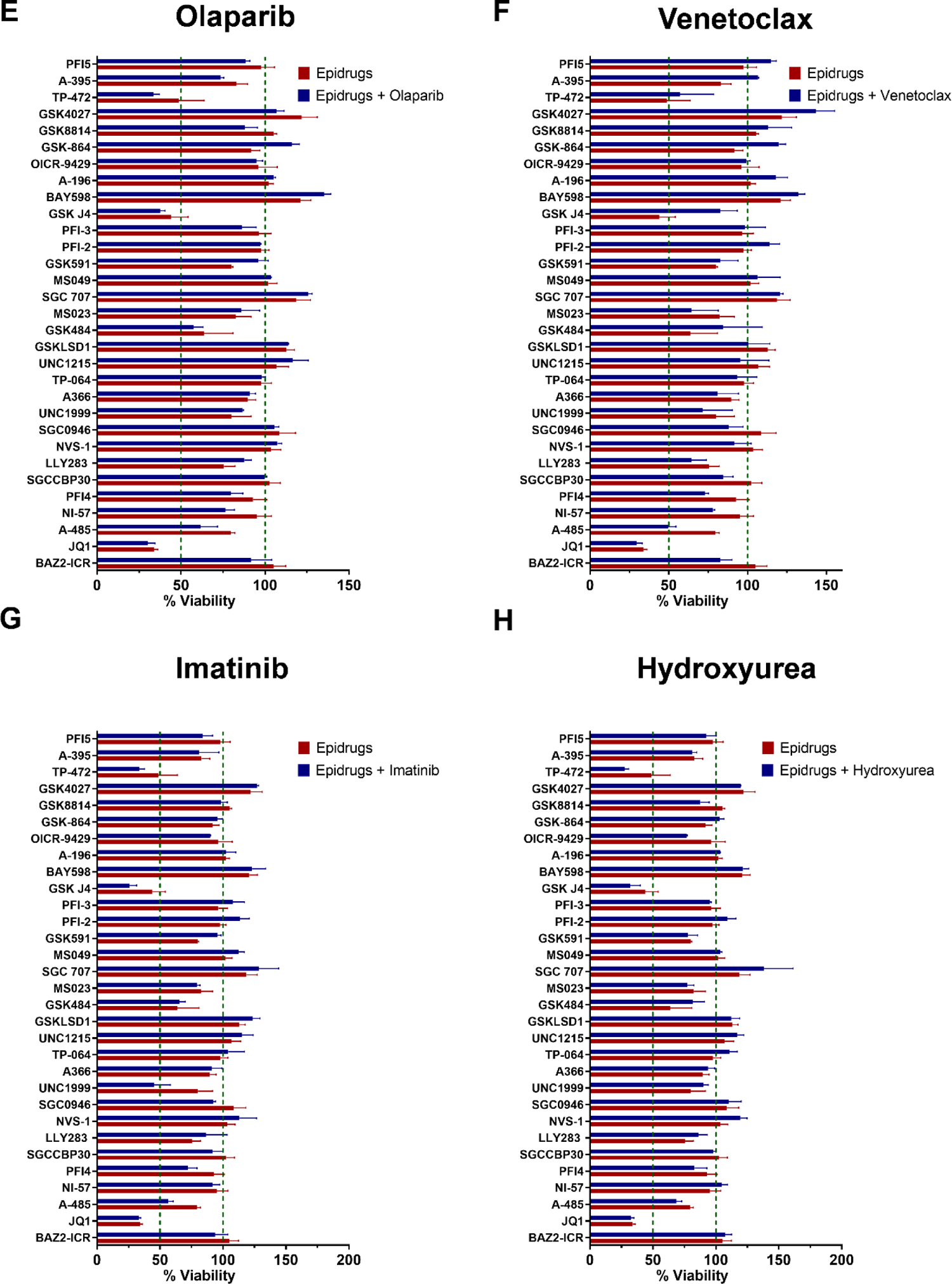
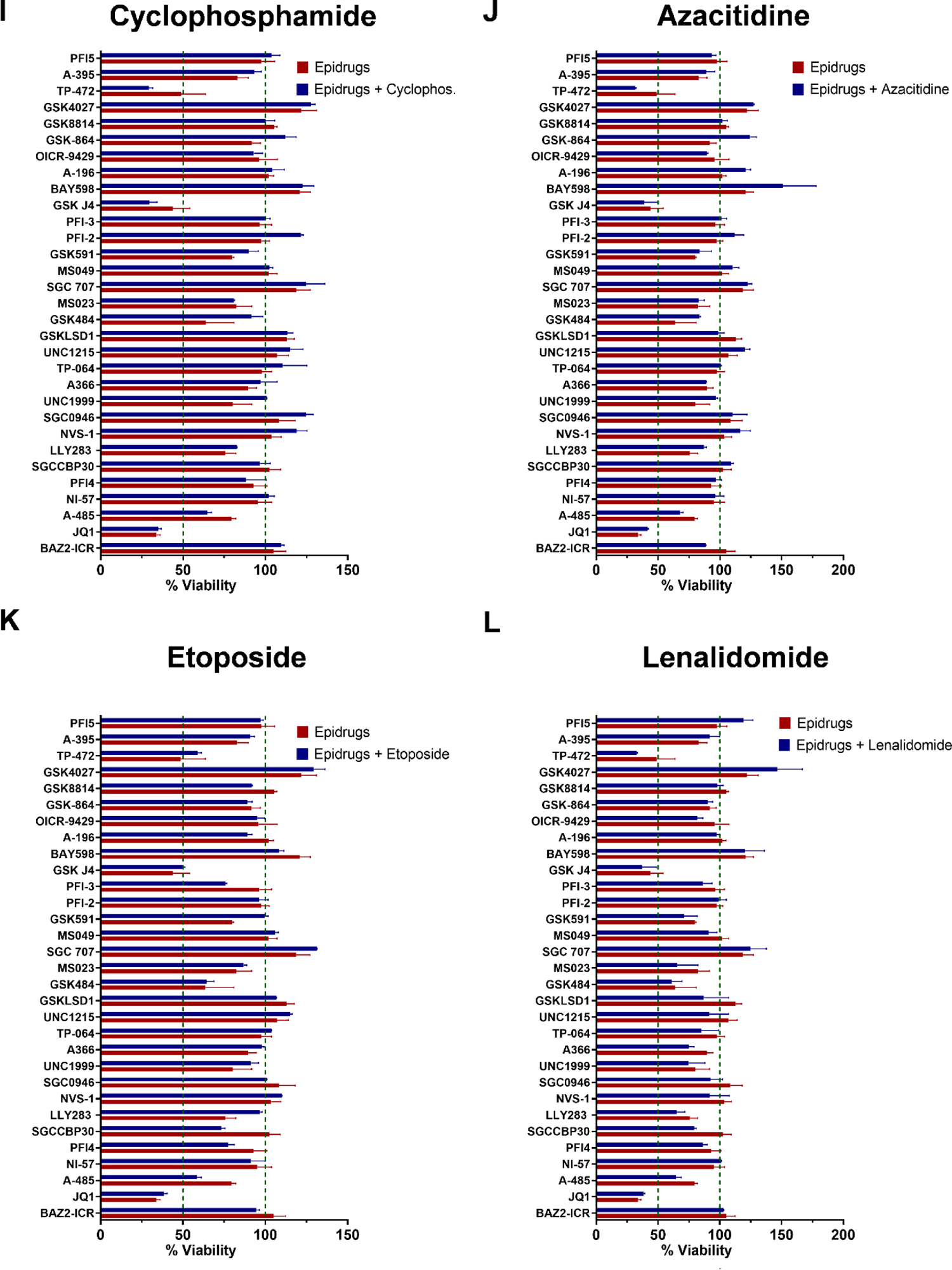
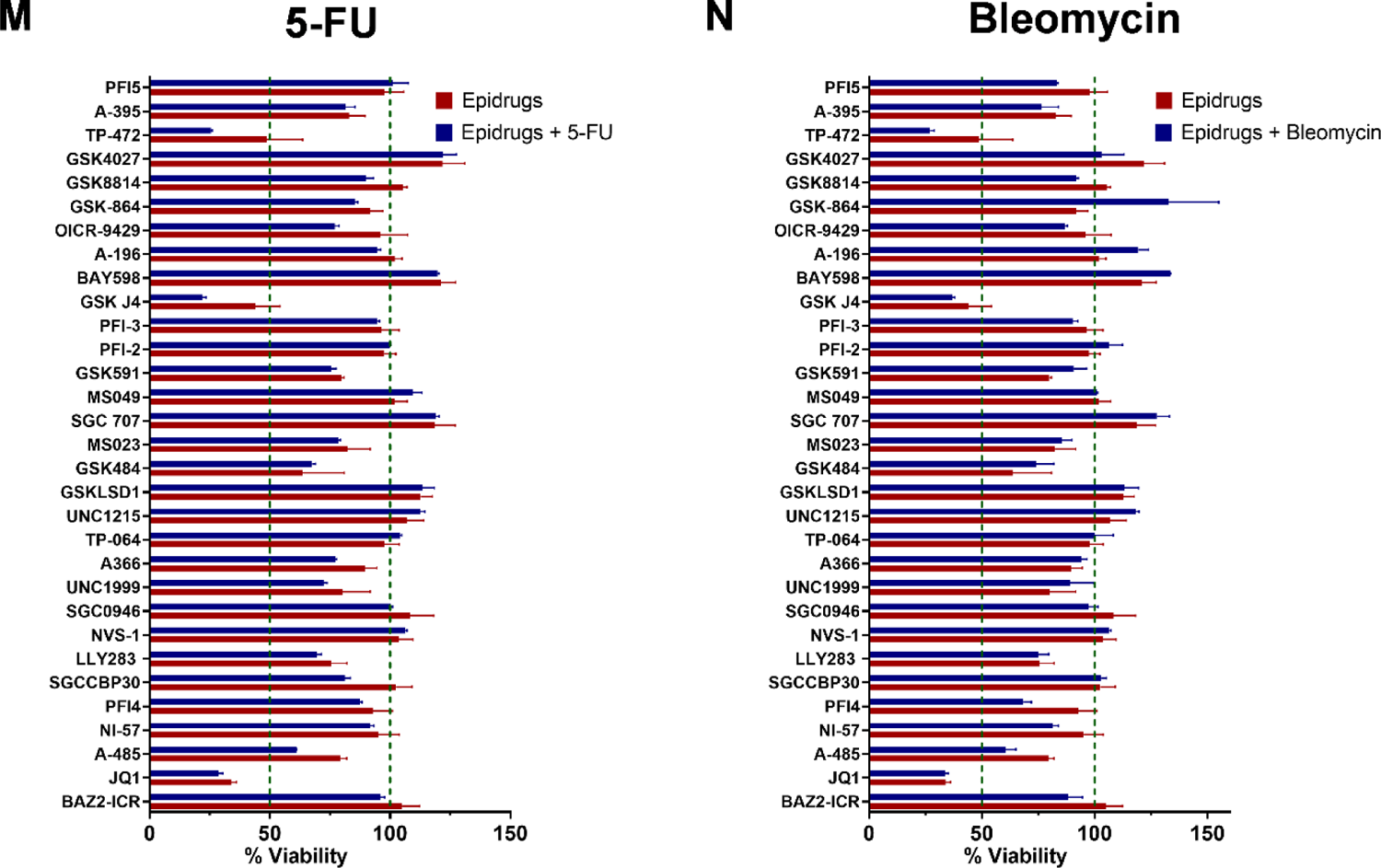
A combinatorial screen identifies epigenetic probes that profoundly potentiate TAK-243 cytotoxicity. Bar charts representing the viability of A549 lung adenocarcinoma cells after treatment with 31 epigenetic probes alone (at concentrations from 1-5 µM) or in combination with 14 anticancer drugs. Viability was assessed by the Resazurin assay after treatment for 72h. These data were used to generate the heatmap in **Fig. 1A**.

**Figure S2.**
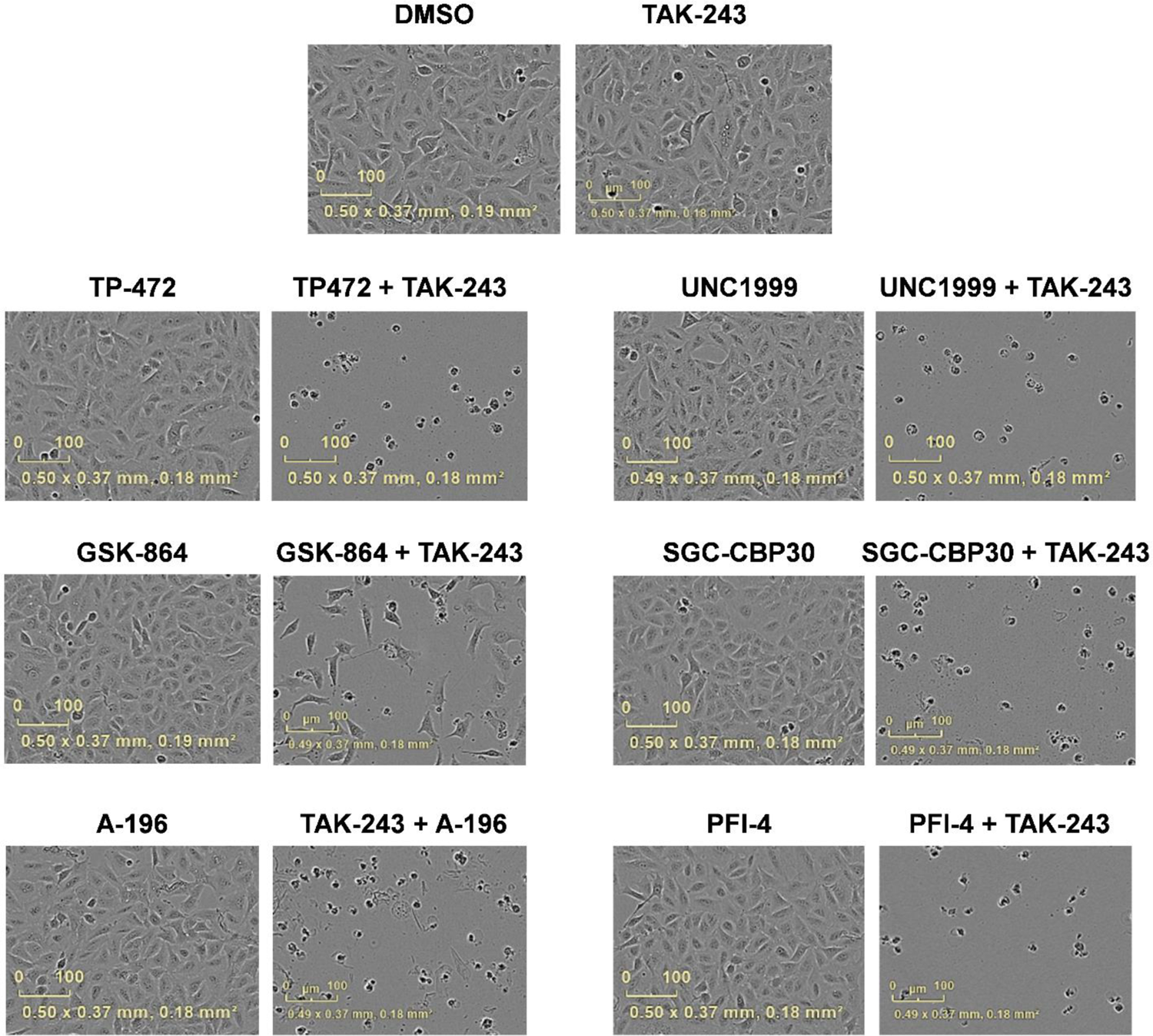
Epigenetic probes potentiate the cytotoxicity of TAK-243 at non-cytotoxic concentrations. Phase contrast images of A549 cells treated with DMSO, TAK-243 (100 nM), epigenetic probes (at concentrations indicated in Table S1), and combinations of TAK-243 and epigenetic probes for 72h. Scale bar = 100 µm.

**Figure S3.**
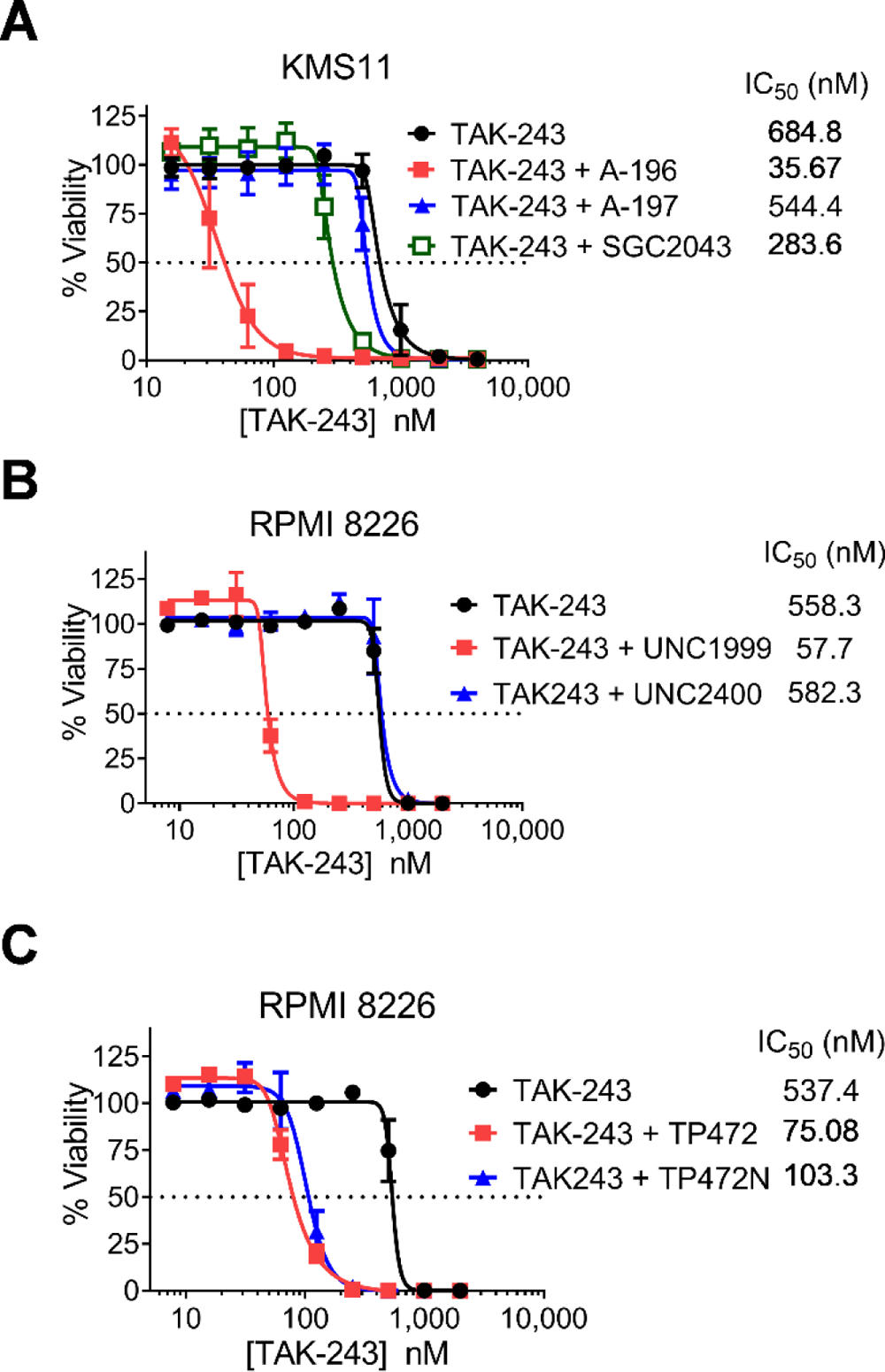
Chemical probes, but not their negative controls, potentiate the cytotoxicity of TAK-243. A) Concentration-response curve of TAK-243 alone or in combination with A-196 (5 µM) or its negative controls A-197 and SGC-2043 (5 µM) in KMS11 myeloma cells. B) Concentration-response curve of TAK-243 alone or in combination with UNC1999 (5 µM) or its negative control UNC2400 (5 µM) in RPMI 8226 myeloma cells C) Concentration-response curve of TAK-243 alone or in combination with TP-472 (5 µM) or its negative control TP-472N (5 µM) in RPMI 8226 myeloma cells. Data points represent the mean ± SEM of at least 3 independent experiments. Insert: median inhibitory concentration (IC_50_) values.

**Figure S4.**
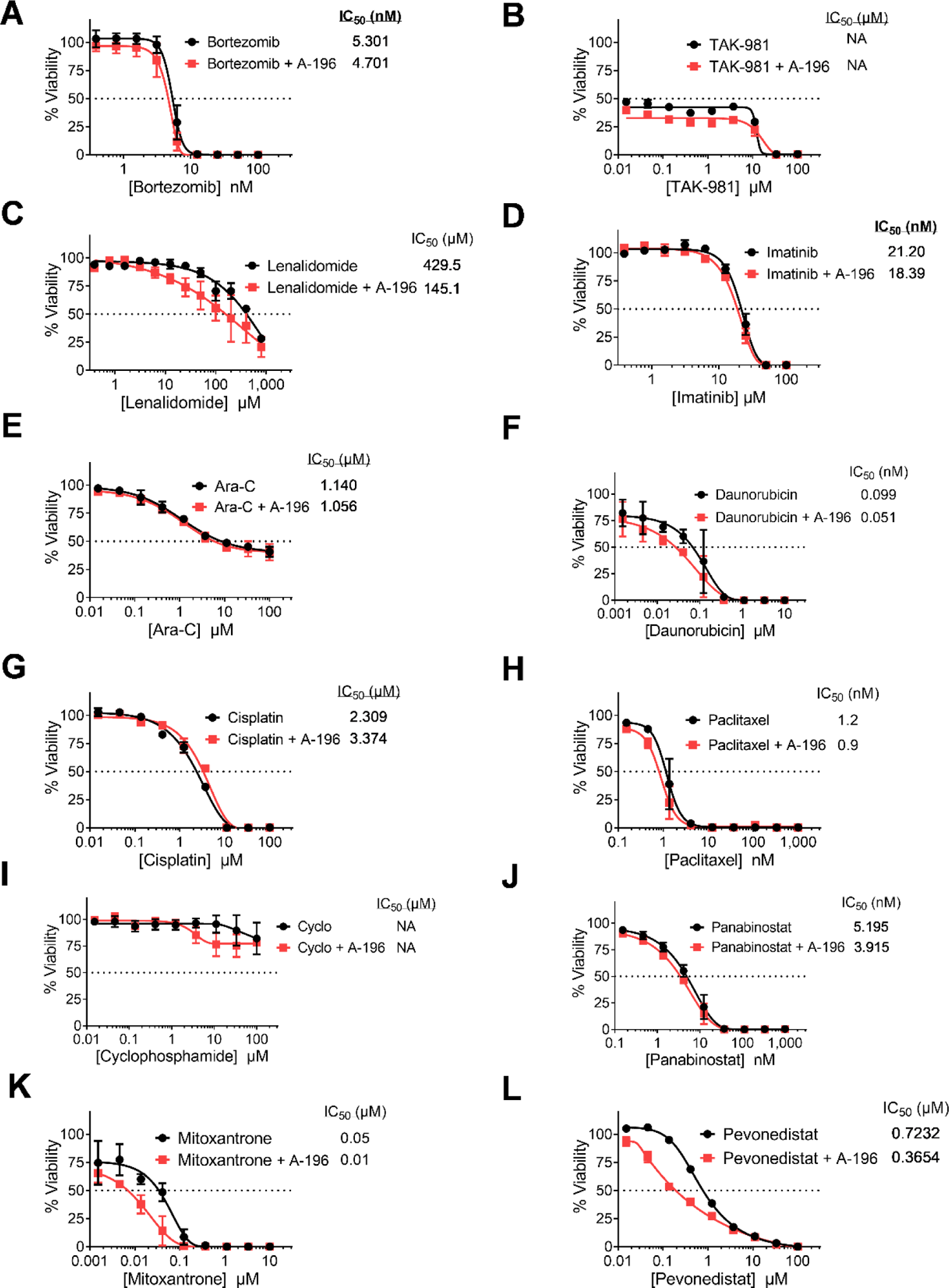
A-196-induced potentiation is relatively selective for TAK-243. A-L) Concentration-response curves of 12 anticancer agents alone or in combination with A-196 (5 µM) in RPMI-8226 cells. Viability was measured by the Resazurin assay after treatment for 72h. Data points represent the mean ± SEM of 2-3 independent experiments. Insert: median inhibitory concentration (IC_50_) values.

**Figure S5.**
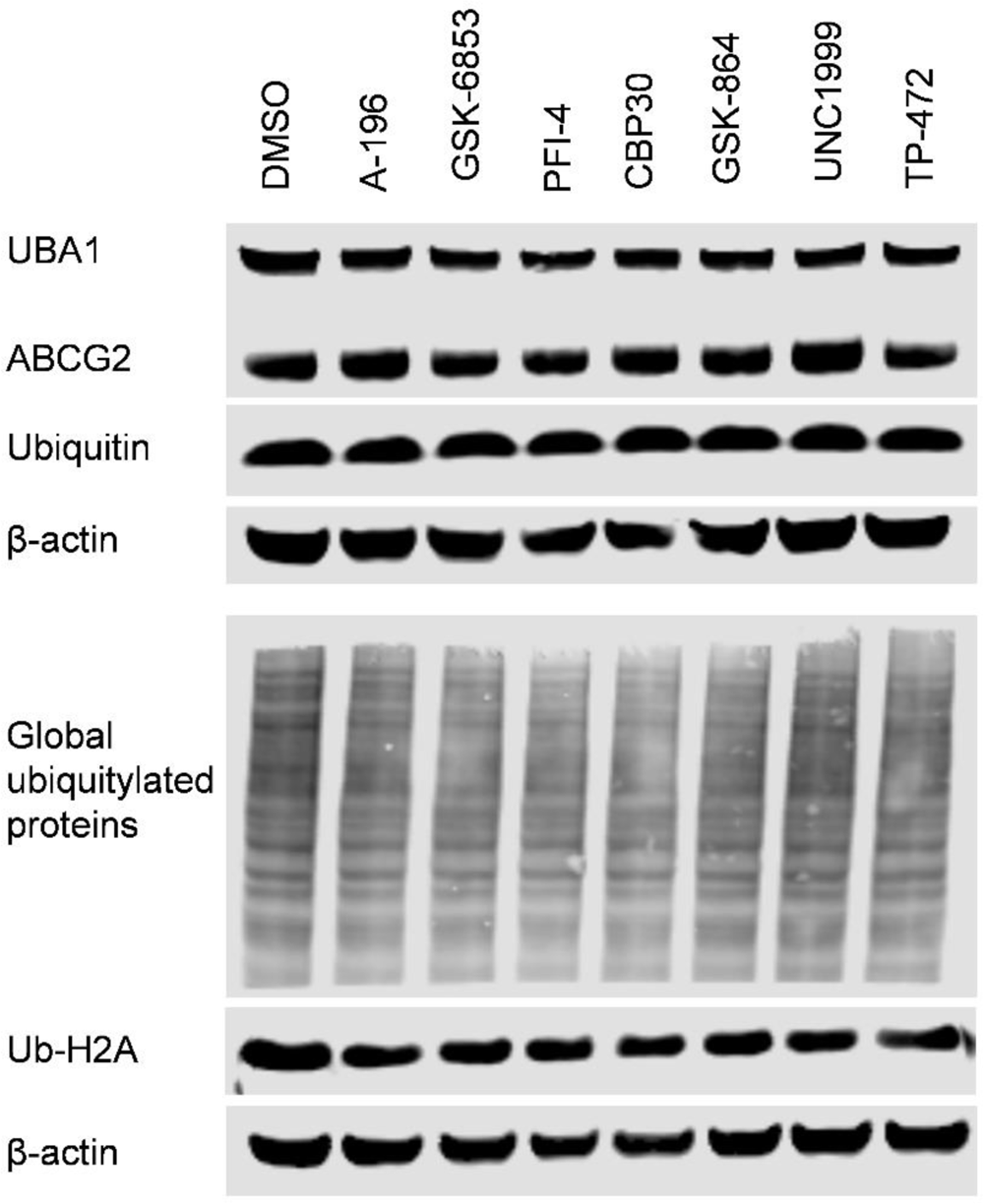
Epigenetic probes do not impact ubiquitylation or ABCG2 expression levels. RPMI 8226 cells were treated with DMSO or epigenetic probes A-196, GSK6853, PFI-4, SGC-CBP30, GSK-864, UNC1999, and TP-472 at 5 µM each for 48h. After treatment, whole cell lysates were prepared and levels of UBA1, ubiquitin, global ubiquitylated proteins, and ubiquitylated H2A (Ub-H2A) were measured by immunoblotting. β-actin was used as a loading control.

**Figure S6.**
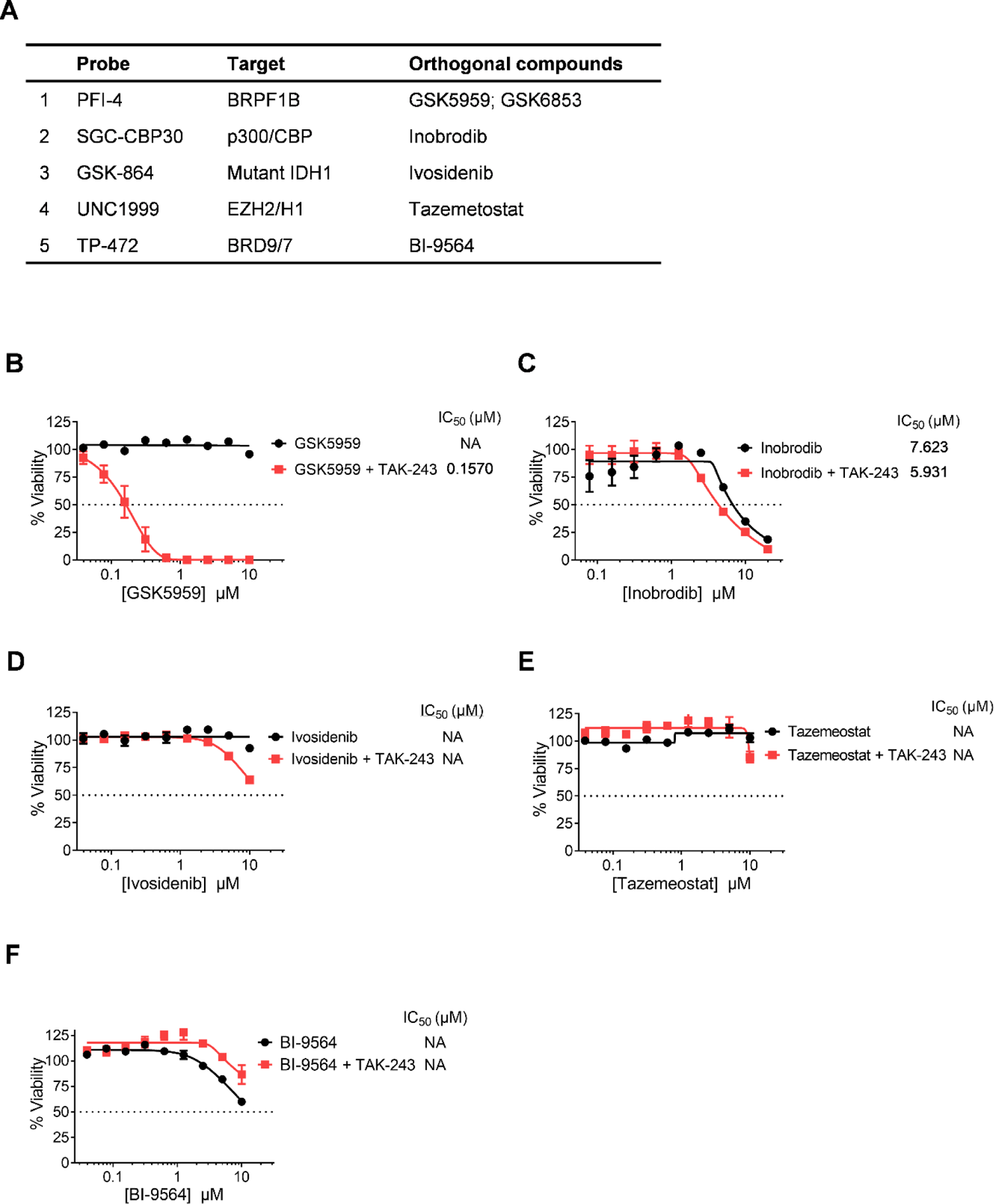
Orthogonal probes/drugs do not phenocopy the potentiation observed with epigenetic probes identified in the screen. A) List of epigenetic probes identified in the screen, their targets, and alternative probes/drugs that target the same epigenetic modulators. B-F) Concentration-response curves of TAK-243 alone or in combination with GSK5959, Inobrodib, Ivosidenib, Tazemetostat, and BI-9564 in RPMI-8226 cells. Data points represent the mean ± SEM of 2-3 independent experiments. Insert: Median inhibitory concentration (IC_50_) values.

**Figure S7.**
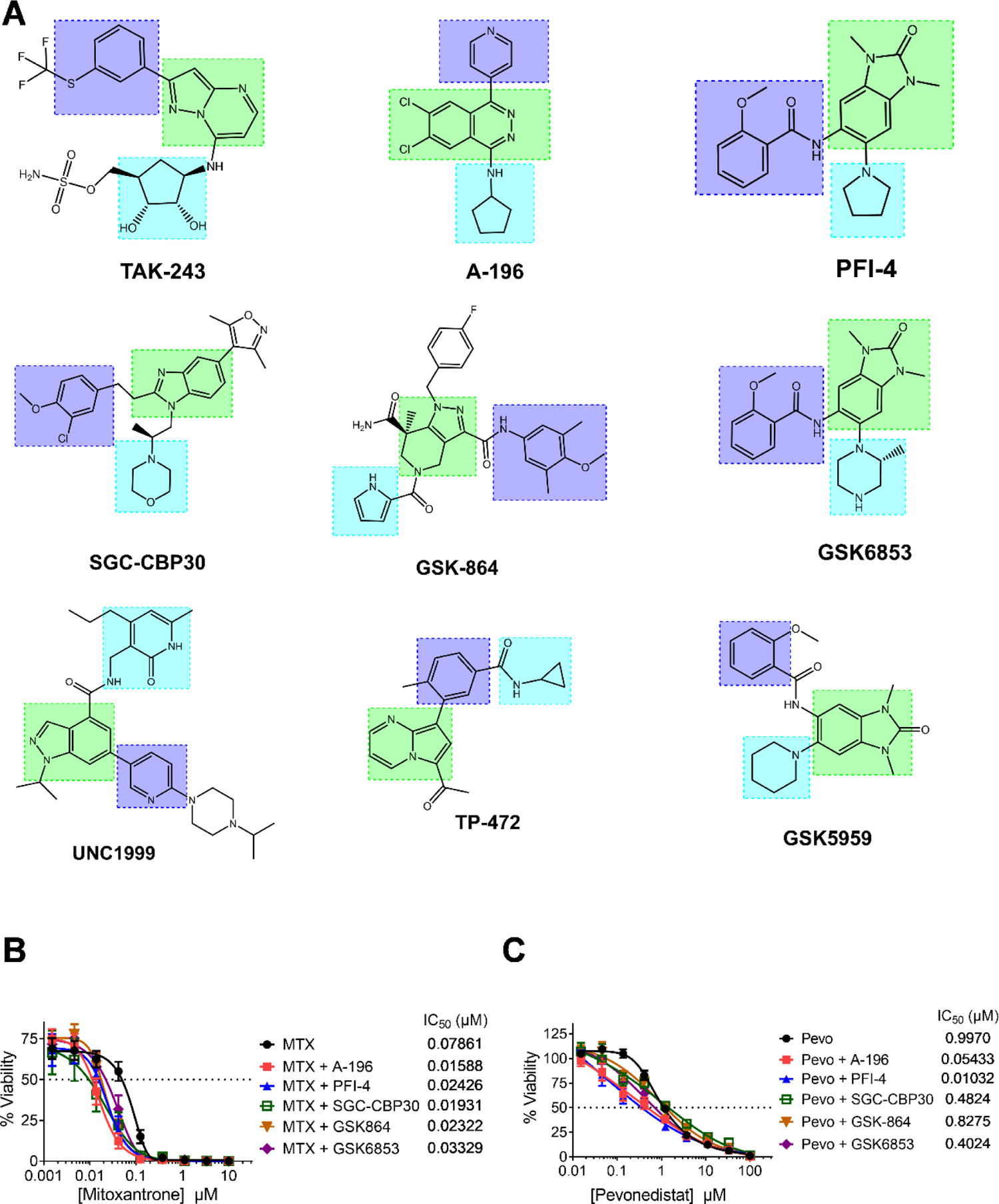
Epigenetic probes share structural similarities with TAK-243 and sensitize cells to the ABCG2 substrates mitoxantrone and pevonedistat. A) Chemical structures of TAK-243, A-196, PFI-4, GSK-CBP30, GSK-864, GSK6853, UNC1999, TP-472, and GSK-5959. Chemically analogous moieties are shaded in similar colors. B) Concentration-response curves of mitoxantrone and pevonedistat after treatment of RPMI 8226 cells with TAK-243 alone and in combination with different epigenetic probes for 72h. Data points represent the mean ± SEM of 3 independent experiments. Insert: Median inhibitory concentration (IC_50_) values.

## References

1. Holohan C, Van Schaeybroeck S, Longley DB, Johnston PG. Cancer drug resistance: an evolving paradigm. Nature reviews. Cancer. Oct 2013;13(10):714–726.

2. Vasan N, Baselga J, Hyman DM. A view on drug resistance in cancer. Nature. Nov 2019;575(7782):299-309.

3. Marine J-C, Dawson S-J, Dawson MA. Non-genetic mechanisms of therapeutic resistance in cancer. Nature Reviews Cancer. 2020;20(12):743–756.

4. Salgia R, Kulkarni P. The Genetic/Non-genetic Duality of Drug ‘Resistance’ in Cancer. Trends in cancer. Feb 2018;4(2):110–118.

5. Dawson MA. The cancer epigenome: Concepts, challenges, and therapeutic opportunities. Science. Mar 17 2017;355(6330):1147-1152.

6. Yuan S, Norgard RJ, Stanger BZ. Cellular Plasticity in Cancer. Cancer Discov. Jul 2019;9(7):837–851.

7. Knoechel B, Roderick JE, Williamson KE, et al. An epigenetic mechanism of resistance to targeted therapy in T cell acute lymphoblastic leukemia. Nature genetics. Apr 2014;46(4):364–370.

8. Hugo W, Shi H, Sun L, et al. Non-genomic and Immune Evolution of Melanoma Acquiring MAPKi Resistance. Cell. Sep 10 2015;162(6):1271–1285.

9. Schenk T, Chen WC, Gollner S, et al. Inhibition of the LSD1 (KDM1A) demethylase reactivates the all-trans-retinoic acid differentiation pathway in acute myeloid leukemia. Nature medicine. Mar 11 2012;18(4):605–611.

10. Shaffer SM, Dunagin MC, Torborg SR, et al. Rare cell variability and drug-induced reprogramming as a mode of cancer drug resistance. Nature. Jun 15 2017;546(7658):431- 435.

11. Wang L, Leite de Oliveira R, Huijberts S, et al. An Acquired Vulnerability of Drug-Resistant Melanoma with Therapeutic Potential. Cell. May 31 2018;173(6):1413–1425 e1414.

12. Morel D, Jeffery D, Aspeslagh S, Almouzni G, Postel-Vinay S. Combining epigenetic drugs with other therapies for solid tumours - past lessons and future promise. Nature reviews. Clinical oncology. Feb 2020;17(2):91–107.

13. Sroczynska P, Cruickshank VA, Bukowski JP, et al. shRNA screening identifies JMJD1C as being required for leukemia maintenance. Blood. Mar 20 2014;123(12):1870–1882.

14. Hoffman GR, Rahal R, Buxton F, et al. Functional epigenetics approach identifies BRM/SMARCA2 as a critical synthetic lethal target in BRG1-deficient cancers. Proceedings of the National Academy of Sciences of the United States of America. Feb 25 2014;111(8):3128–3133.

15. Veschi V, Liu Z, Voss TC, et al. Epigenetic siRNA and Chemical Screens Identify SETD8 Inhibition as a Therapeutic Strategy for p53 Activation in High-Risk Neuroblastoma. Cancer cell. Jan 9 2017;31(1):50–63.

16. Li F, Huang Q, Luster TA, et al. In Vivo Epigenetic CRISPR Screen Identifies Asf1a as an Immunotherapeutic Target in Kras-Mutant Lung Adenocarcinoma. Cancer discovery. Feb 2020;10(2):270–287.

17. Crystal AS, Shaw AT, Sequist LV, et al. Patient-derived models of acquired resistance can identify effective drug combinations for cancer. Science. Dec 19 2014;346(6216):1480- 1486.

18. Shortt J, Ott CJ, Johnstone RW, Bradner JE. A chemical probe toolbox for dissecting the cancer epigenome. Nature reviews. Cancer. Mar 23 2017;17(4):268.

19. Scheer S, Ackloo S, Medina TS, et al. A chemical biology toolbox to study protein methyltransferases and epigenetic signaling. Nat Commun. Jan 3 2019;10(1):19.

20. Wu Q, Heidenreich D, Zhou S, et al. A chemical toolbox for the study of bromodomains and epigenetic signaling. Nature communications. 2019/04/23 2019;10(1):1915.

21. Jafari R, Almqvist H, Axelsson H, et al. The cellular thermal shift assay for evaluating drug target interactions in cells. Nature protocols. 2014;9(9):2100–2122.

22. Barghout SH, Patel PS, Wang X, et al. Preclinical evaluation of the selective small-molecule UBA1 inhibitor, TAK-243, in acute myeloid leukemia. Leukemia. Jan 2019;33(1):37-51.

23. Jackson SM, Manolaridis I, Kowal J, et al. Structural basis of small-molecule inhibition of human multidrug transporter ABCG2. Nature structural & molecular biology. Apr 2018;25(4):333–340.

24. Kowal J, Ni D, Jackson SM, Manolaridis I, Stahlberg H, Locher KP. Structural Basis of Drug Recognition by the Multidrug Transporter ABCG2. Journal of molecular biology. Jun 25 2021;433(13):166980.

25. Yu Q, Ni D, Kowal J, et al. Structures of ABCG2 under turnover conditions reveal a key step in the drug transport mechanism. Nature communications. Jul 19 2021;12(1):4376.

26. Abagyan R, Totrov M, Kuznetsov D. ICM—A new method for protein modeling and design: Applications to docking and structure prediction from the distorted native conformation. Journal of Computational Chemistry. 1994;15(5):488–506.

27. Sastry GM, Adzhigirey M, Day T, Annabhimoju R, Sherman W. Protein and ligand preparation: parameters, protocols, and influence on virtual screening enrichments. Journal of computer-aided molecular design. Mar 2013;27(3):221–234.

28. Halgren TA, Murphy RB, Friesner RA, et al. Glide: a new approach for rapid, accurate docking and scoring. 2. Enrichment factors in database screening. Journal of medicinal chemistry. Mar 25 2004;47(7):1750-1759.

29. Hyer ML, Milhollen MA, Ciavarri J, et al. A small-molecule inhibitor of the ubiquitin activating enzyme for cancer treatment. Nature medicine. Feb 2018;24(2):186–193.

30. Barghout SH, Schimmer AD. E1 Enzymes as Therapeutic Targets in Cancer. Pharmacological Reviews. 2021;73(1):1–58.

31. Barghout SH, Aman A, Nouri K, et al. A genome-wide CRISPR/Cas9 screen in acute myeloid leukemia cells identifies regulators of TAK-243 sensitivity. JCI insight. Mar 8 2021;6(5).

32. Arrowsmith CH, Audia JE, Austin C, et al. The promise and peril of chemical probes. Nature chemical biology. 2015;11(8):536–541.

33. Barghout SH, Schimmer AD. The ubiquitin-activating enzyme, UBA1, as a novel therapeutic target for AML. Oncotarget. 2018;9(76):34198–34199.

34. Toyoda Y, Takada T, Suzuki H. Inhibitors of human ABCG2: from technical background to recent updates with clinical implications. Frontiers in pharmacology. 2019;10:208.

35. Wu Z-X, Yang Y, Wang J-Q, et al. Overexpression of ABCG2 Confers Resistance to MLN7243, a Ubiquitin-Activating Enzyme (UAE) Inhibitor. Frontiers in Cell and Developmental Biology. 2021;9(1727).

36. Manolaridis I, Jackson SM, Taylor NMI, Kowal J, Stahlberg H, Locher KP. Cryo-EM structures of a human ABCG2 mutant trapped in ATP-bound and substrate-bound states. Nature. Nov 2018;563(7731):426-430.

37. Orlando BJ, Liao M. ABCG2 transports anticancer drugs via a closed-to-open switch. Nature communications. May 8 2020;11(1):2264.

38. Gose T, Shafi T, Fukuda Y, et al. ABCG2 requires a single aromatic amino acid to “clamp” substrates and inhibitors into the binding pocket. FASEB journal: official publication of the Federation of American Societies for Experimental Biology. Apr 2020;34(4):4890–4903.

39. Allen JD, van Loevezijn A, Lakhai JM, et al. Potent and specific inhibition of the breast cancer resistance protein multidrug transporter in vitro and in mouse intestine by a novel analogue of fumitremorgin C. Molecular cancer therapeutics. Apr 2002;1(6):417–425.

40. Weidner LD, Zoghbi SS, Lu S, et al. The Inhibitor Ko143 Is Not Specific for ABCG2. The Journal of pharmacology and experimental therapeutics. Sep 2015;354(3):384–393.

41. Dantzig AH, Shepard RL, Law KL, et al. Selectivity of the multidrug resistance modulator, LY335979, for P-glycoprotein and effect on cytochrome P-450 activities. The Journal of pharmacology and experimental therapeutics. Aug 1999;290(2):854-862.

42. Wimalasena VK, Wang T, Sigua LH, Durbin AD, Qi J. Using Chemical Epigenetics to Target Cancer. Mol Cell. May 13 2020.

43. Setton J, Zinda M, Riaz N, et al. Synthetic Lethality in Cancer Therapeutics: The Next Generation. Cancer discovery. Jul 2021;11(7):1626–1635.

44. Liu G, Yu J, Wu R, et al. GRP78 determines glioblastoma sensitivity to UBA1 inhibition-induced UPR signaling and cell death. Cell death & disease. Jul 23 2021;12(8):733.

45. Elias R, Tcheuyap VT, Kaushik AK, et al. A renal cell carcinoma tumorgraft platform to advance precision medicine. Cell reports. Nov 23 2021;37(8):110055.

46. Majeed S, Aparnathi MK, Nixon KCJ, et al. Targeting the ubiquitin-proteasome system using the UBA1 inhibitor TAK-243 is a potential therapeutic strategy for small cell lung cancer. Clinical cancer research: an official journal of the American Association for Cancer Research. Feb 14 2022.

47. Zhuang J, Shirazi F, Singh RK, et al. Ubiquitin-activating enzyme inhibition induces an unfolded protein response and overcomes drug resistance in myeloma. Blood. 2019;133(14):1572–1584.

48. Best S, Hashiguchi T, Kittai A, et al. Targeting ubiquitin-activating enzyme induces ER stress-mediated apoptosis in B-cell lymphoma cells. Blood Adv. 2019;3(1):51–62.

49. Best S, Liu T, Bruss N, Kittai A, Berger A, Danilov AV. Pharmacologic inhibition of the ubiquitin-activating enzyme induces ER stress and apoptosis in chronic lymphocytic leukemia and ibrutinib-resistant mantle cell lymphoma cells. Leukemia & Lymphoma. 2019/10/15 2019;60(12):2946-2950.

50. Lee J, Schapira M. The Promise and Peril of Chemical Probe Negative Controls. ACS chemical biology. Apr 16 2021;16(4):579–585.

51. Murai Y, Jo U, Murai J, et al. SLFN11 Inactivation Induces Proteotoxic Stress and Sensitizes Cancer Cells to Ubiquitin Activating Enzyme Inhibitor TAK-243. Cancer Res. 2021;81(11):3067–3078.

52. Miyake K, Mickley L, Litman T, et al. Molecular cloning of cDNAs which are highly overexpressed in mitoxantrone-resistant cells: demonstration of homology to ABC transport genes. Cancer Res. Jan 1 1999;59(1):8–13.

53. Doyle LA, Yang W, Abruzzo LV, et al. A multidrug resistance transporter from human MCF-7 breast cancer cells. Proc Natl Acad Sci U S A. Dec 22 1998;95(26):15665–15670.

54. Wu Z-X, Mai Q, Yang Y, et al. Overexpression of human ATP-binding cassette transporter ABCG2 contributes to reducing the cytotoxicity of GSK1070916 in cancer cells. Biomedicine & Pharmacotherapy. 2021;136:111223.

## References

1. Taylor AP, Swewczyk M, Kennedy S, et al. Selective, Small-Molecule Co-Factor Binding Site Inhibition of a Su(var)3-9, Enhancer of Zeste, Trithorax Domain Containing Lysine Methyltransferase. Journal of medicinal chemistry. Sep 12 2019;62(17):7669–7683.

2. He Y, Selvaraju S, Curtin ML, et al. The EED protein-protein interaction inhibitor A-395 inactivates the PRC2 complex. Nature chemical biology. Apr 2017;13(4):389–395.

3. Mason LD, Chava S, Reddi KK, Gupta R. The BRD9/7 Inhibitor TP-472 Blocks Melanoma Tumor Growth by Suppressing ECM-Mediated Oncogenic Signaling and Inducing Apoptosis. Cancers. Nov 3 2021;13(21).

4. Humphreys PG, Bamborough P, Chung CW, et al. Discovery of a Potent, Cell Penetrant, and Selective p300/CBP-Associated Factor (PCAF)/General Control Nonderepressible 5 (GCN5) Bromodomain Chemical Probe. Journal of medicinal chemistry. Jan 26 2017;60(2):695-709.

5. Bamborough P, Chung CW, Demont EH, et al. A Chemical Probe for the ATAD2 Bromodomain. Angewandte Chemie. Sep 12 2016;55(38):11382–11386.

6. Okoye-Okafor UC, Bartholdy B, Cartier J, et al. New IDH1 mutant inhibitors for treatment of acute myeloid leukemia. Nature chemical biology. Nov 2015;11(11):878–886.

7. Grebien F, Vedadi M, Getlik M, et al. Pharmacological targeting of the Wdr5-MLL interaction in C/EBPalpha N-terminal leukemia. Nature chemical biology. Aug 2015;11(8):571–578.

8. Bromberg KD, Mitchell TR, Upadhyay AK, et al. The SUV4-20 inhibitor A-196 verifies a role for epigenetics in genomic integrity. Nature chemical biology. Mar 2017;13(3):317–324.

9. Eggert E, Hillig RC, Koehr S, et al. Discovery and Characterization of a Highly Potent and Selective Aminopyrazoline-Based in Vivo Probe (BAY-598) for the Protein Lysine Methyltransferase SMYD2. Journal of medicinal chemistry. May 26 2016;59(10):4578–4600.

10. Kruidenier L, Chung CW, Cheng Z, et al. A selective jumonji H3K27 demethylase inhibitor modulates the proinflammatory macrophage response. Nature. Aug 16 2012;488(7411):404-408.

11. Fedorov O, Castex J, Tallant C, et al. Selective targeting of the BRG/PB1 bromodomains impairs embryonic and trophoblast stem cell maintenance. Science advances. Nov 2015;1(10):e1500723.

12. Barsyte-Lovejoy D, Li F, Oudhoff MJ, et al. (R)-PFI-2 is a potent and selective inhibitor of SETD7 methyltransferase activity in cells. Proceedings of the National Academy of Sciences of the United States of America. Sep 2 2014;111(35):12853–12858.

13. Fong JY, Pignata L, Goy PA, et al. Therapeutic Targeting of RNA Splicing Catalysis through Inhibition of Protein Arginine Methylation. Cancer cell. Aug 12 2019;36(2):194–209 e199.

14. Shen Y, Szewczyk MM, Eram MS, et al. Discovery of a Potent, Selective, and Cell-Active Dual Inhibitor of Protein Arginine Methyltransferase 4 and Protein Arginine Methyltransferase 6. Journal of medicinal chemistry. Oct 13 2016;59(19):9124-9139.

15. Kaniskan HU, Szewczyk MM, Yu Z, et al. A potent, selective and cell-active allosteric inhibitor of protein arginine methyltransferase 3 (PRMT3). Angewandte Chemie. Apr 20 2015;54(17):5166–5170.

16. Eram MS, Shen Y, Szewczyk M, et al. A Potent, Selective, and Cell-Active Inhibitor of Human Type I Protein Arginine Methyltransferases. ACS chemical biology. Mar 18 2016;11(3):772–781.

17. Lewis HD, Liddle J, Coote JE, et al. Inhibition of PAD4 activity is sufficient to disrupt mouse and human NET formation. Nature chemical biology. Mar 2015;11(3):189–191.

18. Cusan M, Cai SF, Mohammad HP, et al. LSD1 inhibition exerts its antileukemic effect by recommissioning PU.1- and C/EBPalpha-dependent enhancers in AML. Blood. Apr 12 2018;131(15):1730-1742.

19. James LI, Barsyte-Lovejoy D, Zhong N, et al. Discovery of a chemical probe for the L3MBTL3 methyllysine reader domain. Nature chemical biology. Mar 2013;9(3):184–191.

20. Nakayama K, Szewczyk MM, Dela Sena C, et al. TP-064, a potent and selective small molecule inhibitor of PRMT4 for multiple myeloma. Oncotarget. Apr 6 2018;9(26):18480–18493.

21. Sweis RF, Pliushchev M, Brown PJ, et al. Discovery and development of potent and selective inhibitors of histone methyltransferase g9a. ACS medicinal chemistry letters. Feb 13 2014;5(2):205–209.

22. Konze KD, Ma A, Li F, et al. An orally bioavailable chemical probe of the Lysine Methyltransferases EZH2 and EZH1. ACS chemical biology. 2013;8(6):1324–1334.

23. Yu W, Chory EJ, Wernimont AK, et al. Catalytic site remodelling of the DOT1L methyltransferase by selective inhibitors. Nature communications. 2012;3:1288.

24. Park SG, Lee D, Seo HR, Lee SA, Kwon J. Cytotoxic activity of bromodomain inhibitor NVS-CECR2-1 on human cancer cells. Scientific reports. Oct 1 2020;10(1):16330.

25. Bonday ZQ, Cortez GS, Grogan MJ, et al. LLY-283, a Potent and Selective Inhibitor of Arginine Methyltransferase 5, PRMT5, with Antitumor Activity. ACS medicinal chemistry letters. Jul 12 2018;9(7):612-617.

26. Hammitzsch A, Tallant C, Fedorov O, et al. CBP30, a selective CBP/p300 bromodomain inhibitor, suppresses human Th17 responses. Proceedings of the National Academy of Sciences of the United States of America. Aug 25 2015;112(34):10768–10773.

27. Meier JC, Tallant C, Fedorov O, et al. Selective Targeting of Bromodomains of the Bromodomain-PHD Fingers Family Impairs Osteoclast Differentiation. ACS chemical biology. Oct 20 2017;12(10):2619–2630.

28. Igoe N, Bayle ED, Tallant C, et al. Design of a Chemical Probe for the Bromodomain and Plant Homeodomain Finger-Containing (BRPF) Family of Proteins. Journal of medicinal chemistry. Aug 24 2017;60(16):6998–7011.

29. Lasko LM, Jakob CG, Edalji RP, et al. Discovery of a selective catalytic p300/CBP inhibitor that targets lineage-specific tumours. Nature. Oct 5 2017;550(7674):128-132.

30. Filippakopoulos P, Qi J, Picaud S, et al. Selective inhibition of BET bromodomains. Nature. Dec 23 2010;468(7327):1067-1073.

31. Drouin L, McGrath S, Vidler LR, et al. Structure enabled design of BAZ2-ICR, a chemical probe targeting the bromodomains of BAZ2A and BAZ2B. Journal of medicinal chemistry. Mar 12 2015;58(5):2553–2559.

32. Hyer ML, Milhollen MA, Ciavarri J, et al. A small-molecule inhibitor of the ubiquitin activating enzyme for cancer treatment. Nature medicine. Feb 2018;24(2):186–193.

33. Soucy TA, Smith PG, Milhollen MA, et al. An inhibitor of NEDD8-activating enzyme as a new approach to treat cancer. Nature. Apr 9 2009;458(7239):732-736.

34. Lightcap ES, Yu P, Grossman S, et al. A small-molecule SUMOylation inhibitor activates antitumor immune responses and potentiates immune therapies in preclinical models. Science translational medicine. Sep 15 2021;13(611):eaba7791.

35. Rottenberg S, Disler C, Perego P. The rediscovery of platinum-based cancer therapy. Nature reviews. Cancer. Jan 2021;21(1):37–50.

36. Curtin NJ, Szabo C. Poly(ADP-ribose) polymerase inhibition: past, present and future. Nature reviews. Drug discovery. Oct 2020;19(10):711–736.

37. Ashkenazi A, Fairbrother WJ, Leverson JD, Souers AJ. From basic apoptosis discoveries to advanced selective BCL-2 family inhibitors. Nature reviews. Drug discovery. Apr 2017;16(4):273–284.

38. Cohen P, Cross D, Janne PA. Kinase drug discovery 20 years after imatinib: progress and future directions. Nature reviews. Drug discovery. Jul 2021;20(7):551–569.

39. Musialek MW, Rybaczek D. Hydroxyurea-The Good, the Bad and the Ugly. Genes. Jul 19 2021;12(7).

40. Emadi A, Jones RJ, Brodsky RA. Cyclophosphamide and cancer: golden anniversary. Nature reviews. Clinical oncology. Nov 2009;6(11):638–647.

41. Issa JP, Kantarjian HM, Kirkpatrick P. Azacitidine. Nature reviews. Drug discovery. Apr 2005;4(4):275–276.

42. Zhang W, Gou P, Dupret JM, Chomienne C, Rodrigues-Lima F. Etoposide, an anticancer drug involved in therapy-related secondary leukemia: Enzymes at play. Translational oncology. Oct 2021;14(10):101169.

43. Jan M, Sperling AS, Ebert BL. Cancer therapies based on targeted protein degradation - lessons learned with lenalidomide. Nature reviews. Clinical oncology. Jul 2021;18(7):401–417.

44. Vodenkova S, Buchler T, Cervena K, Veskrnova V, Vodicka P, Vymetalkova V. 5-fluorouracil and other fluoropyrimidines in colorectal cancer: Past, present and future. Pharmacology & therapeutics. Feb 2020;206:107447.

45. Brandt JP, Gerriets V. Bleomycin. StatPearls. Treasure Island (FL)2022.

